# Improved Estimation of Correlation Accuracy for Machine Learning Brain-Phenotype Associations

**DOI:** 10.1101/2025.11.26.690778

**Authors:** Megan T. Jones, Ishaan Gadiyar, Xinyu Zhang, Kaidi Kang, Jinyuan Liu, Andrew Chen, Jakob Seidlitz, Aaron Alexander-Bloch, Edward Kennedy, Simon Vandekar

**Affiliations:** Vanderbilt University; Wake Forest University; Vanderbilt University Medical Center; Medical University of South Carolina; University of Pennsylvania; Children’s Hospital of Philadelphia; Carnegie Mellon University

## Abstract

Machine learning is used in neuroscience to examine brain-phenotype associations and facilitate individual prediction from high-dimensional brain imaging. For continuous phenotypes, Pearson’s correlation between the observed and predicted phenotype is used to quantify model accuracy in testing data. However, recent research suggests millions of samples may be needed to reliably estimate the maximum achievable predictive accuracy (MAPA). We formally define the MAPA and show that Pearson’s estimator is biased for this quantity and its confidence intervals fail to capture the target. We develop a semiparametric (double machine learning) one-step estimator that more accurately estimates the MAPA and yields valid confidence intervals across flexible machine learning settings. Analyzing data from the Reproducible Brain Charts dataset, we show that this estimator has smaller bias when estimating brain-phenotype associations of neuroimaging data with age and psychopathology phenotypes. We show that MAPA for psychopathology factor scores using machine learning models built on structural and functional imaging measures is not better than using demographic and nuisance covariates alone.

## Introduction

Brain-phenotype association studies examine the relationship between brain measurements obtained using magnetic resonance imaging (MRI) and individual differences in psychiatric, cognitive, or behavioral phenotypes^1^. Machine learning (ML) brain-phenotype associations use high-dimensional brain imaging features simultaneously to predict phenotypes, often to evaluate complex associations between brain and behavior. This is distinct from mass-univariate methods, where each brain feature is modeled as the outcome. There is evidence that ML associations yield bigger effect sizes and better replicability^1–3^, an important consideration given recent evidence of the relatively low magnitude of most brain-behavior associations^1^. With increasing sophistication of artificial intelligence and deep learning image analysis and increasing sample sizes, the utility of ML methods to evaluate brain-phenotype associations is likely to grow^1,4–8^.

An essential feature of ML brain-phenotype efforts is a report of the overall predictive accuracy of the ML model. Prediction accuracy provides an interpretable estimate of the utility of the brain imaging feature to identify values of the phenotype. With research on neuroimaging brain-phenotype associations still in a relatively early stage, such indices help identify which brain measures may be useful biomarkers of behavior or clinical conditions^7,9^. For continuous phenotypes, predictive accuracy is routinely estimated using Pearson’s correlation estimator in held-out data not used to train the model^7,10^.

Recent work has demonstrated a sample-size-dependent negative bias of the standard Pearson correlation estimator, suggesting the number of samples needed for an unbiased estimate varies by brain-phenotype association, sometimes requiring more than one million samples^2,9^. This bias makes it difficult to determine if a model’s low predictive accuracy is due to limited sample size or a truly uninformative association, and suggests it is infeasible to accurately evaluate prediction models in typical studies^9^. In fact, these accuracy estimates are specific to the particular estimated model and sample size from the training dataset. Thus, they cannot be interpreted as a measure of the maximum achievable predictive accuracy (MAPA) we can hope to obtain for a given brain-phenotype association and ML pipeline in the population.

Here, we formally define MAPA for a given ML model class, brain measure, and phenotype, and show that Pearson’s correlation fails to capture this target value. We use modern semiparametric statistical theory to modify Pearson’s correlation estimator to improve replicability and validity as an index of brain-phenotype association strength. This new estimator allows neuroscientists to compare the prediction accuracy of multiple ML models and estimate large-sample valid confidence intervals (CIs). We use the approach to compare MAPA for age and psychopathology factor scores using a base model including non-imaging and nuisance features with models including structural and functional neuroimaging features. Our results suggest neuroimaging features do not increase MAPA for psychopathology over the base model. The one-step estimator (available for R via https://github.com/statimagcoll/semicor) is model-agnostic, able to be used with a broad range of ML algorithms and provides a computationally efficient estimate and CI procedure to improve accuracy in reporting ML brain-phenotype associations.

## Results

### Dataset details

The Reproducible Brain Charts (RBC) project process raw neuroimaging data using sMRIPrep, FreeSurfer, and the C-PAC pipeline^11,12^, and harmonized psychiatric phenotypes (general psychopathology “P-factor”, externalizing, internalizing, and attention-hyperactivity symptoms) are derived from a factor analysis^13,14^ (**Methods**; **Table 1**).

**Table 1.** Participant characteristics in the Reproducible Brain Charts (RBC) dataset. Brain-phenotype associations are examined for 3074 participants across the 5 studies included in the RBC. Demographic factors include age and sex. Psychopathologic factors include P-factor, internalizing score, and externalizing score. Imaging-related quality measures include mean framewise displacement for functional data and Euler number for structural data.

|  | Overall<br>(n=3074) |
| --- | --- |
| **Age (years)** |  |
| Mean (SD) | 13.1 (3.85) |
| Median [Min, Max] | 12.8 [5.02, 23.0] |
| **Sex** |  |
| Female | 1443 (46.9%) |
| Male | 1631 (53.1%) |
| **Internalizing Score** |  |
| Mean (SD) | -0.0120 (0.996) |
| Median [Min, Max] | -0.150 [-2.31, 2.82] |
| **Externalizing Score** |  |
| Mean (SD) | -0.112 (0.898) |
| Median [Min, Max] | -0.314 [-2.22, 3.89] |
| **P-Factor** |  |
| Mean (SD) | -0.145 (1.00) |
| Median [Min, Max] | -0.194 [-1.61, 2.77] |
| **Mean Framewise Displacement** |  |
| Mean (SD) | 0.106 (0.104) |
| Median [Min, Max] | 0.0756 [0.0140, 1.75] |
| **Euler Number** |  |
| Mean (SD) | -107 (67.9) |
| Median [Min, Max] | -92.0 [-1490, -11.0] |

### ML brain-phenotype associations should target MAPA

We use 500 realistic plasmode simulations to emphasize that ML brain-phenotype associations should estimate the true MAPA for a given class of ML models^15^ (**Figure 1A**). Our simulations resample data of 3486 individuals (**Table 1**) from the RBC to distinguish three types of correlation accuracy indices predicting age from regional gray matter volume (GMV) within 62 regions of the Desikan-Killiany-Tourville (DKT) atlas^15–17^. A build dataset of 697 individuals is used to fit a true ML model using a random forest. We resample the remaining 2789 individuals’ MRI data, with replacement, for sample sizes ranging from 100 to 20,000 and add uniform error to the predictions from the random forest model to generate unique phenotype values. We use 5-fold cross-fitting with nested 10-fold cross-validated ridge regression (for tuning) as our restricted model class (**Methods**).

**Figure 1.**
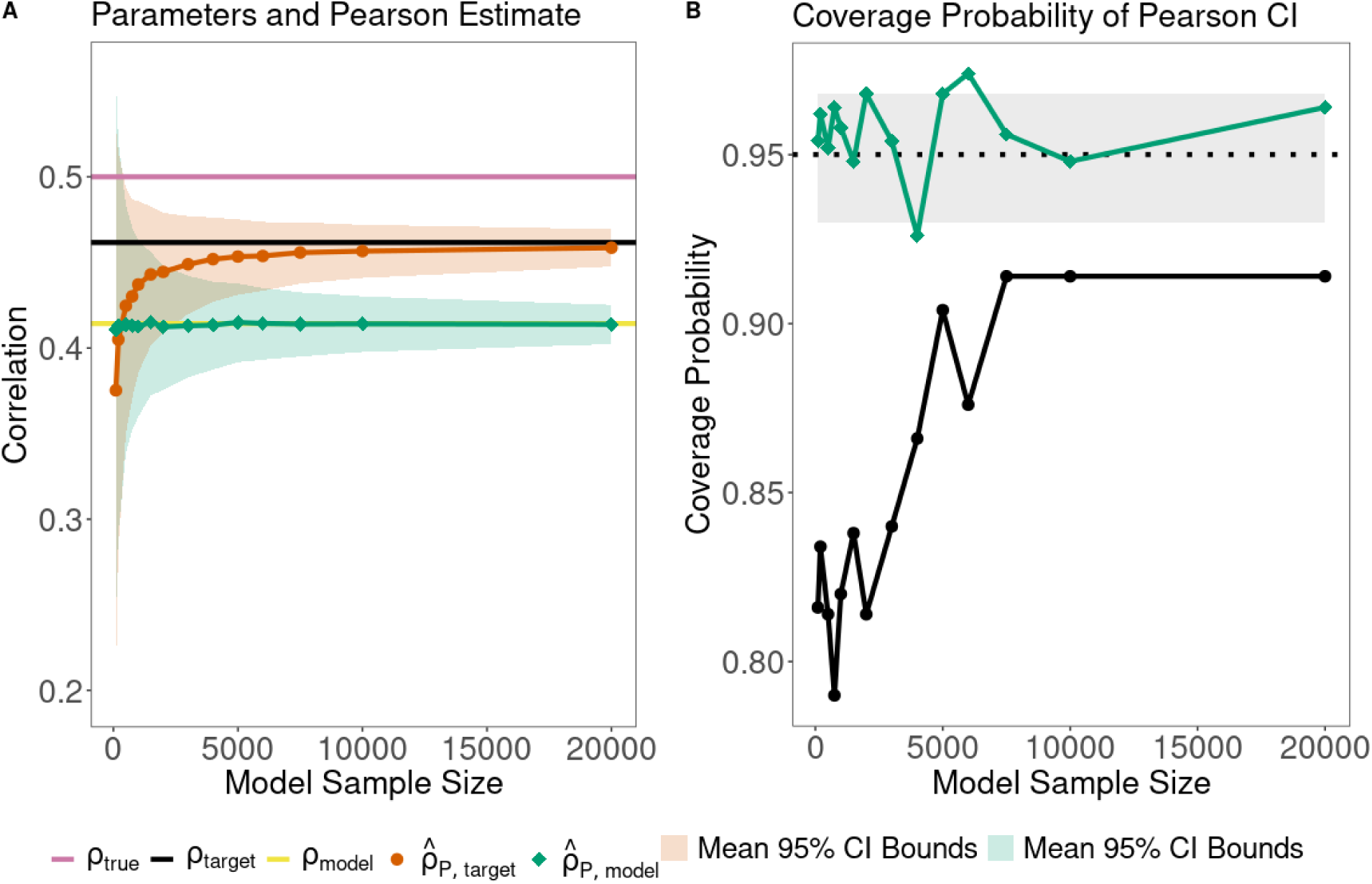
Pearson’s correlation estimator performs poorly for machine learning associations with less than 10,000 samples when estimating prediction accuracy of brain volumes for age. **A)** Simulation results demonstrating the statistical concepts: MPA (pink) denotes the true correlation between the imaging data and non-imaging phenotype, which is the best prediction accuracy possible for the phenotype. MAPA (black) denotes the best possible prediction accuracy of models under consideration (e.g. all possible ridge regression models). TPA (yellow) represents the true prediction accuracy for a particular model that is fit to a specific training set of *n* observations (*n* = 500 here). The orange and green lines indicate the expected value of the Pearson estimator for estimating MAPA and TPA, respectively. Average 95% confidence bounds are shaded. With a more flexible model class (e.g. random forest), the black line will move closer to the pink line. The Pearson estimator is biased downward for estimating MAPA and is unbiased for estimating TPA. **B)** Confidence interval accuracy (coverage probability) of Pearson’s correlation estimator for estimating MAPA (orange) and TPA (green). A good confidence interval should have coverage in the gray region. Coverage is in the expected range given 500 simulations for TPA, but below the nominal level for MAPA, even at 20,000 observations.

We identify three parameters (**Figure 1A**): Maximum predictive accuracy (MPA) quantifies the correlation between the phenotype and the arbitrarily complex, unknown true conditional expectation of the phenotype, given the input features. This represents the highest possible predictive accuracy from the infinite set of all possible predictive models using the selected brain features. Because the true conditional expectation is not observable, MPA cannot be measured directly. We therefore focus on MAPA, which quantifies the maximum predictive accuracy *for a given class of ML models.* In this simulated example, the phenotype is generated from a random forest model so that MPA = 0.5. MAPA is the maximum predictive accuracy within the class of ridge regression models fit to the same features; in the simulated example, MAPA = 0.46. The difference between MPA and MAPA is approximation bias due to the true model being outside the ML model class^18,19^. Considering a more flexible model class (e.g., random forest models) will reduce the approximation bias by moving MAPA closer to MPA.

The third parameter, TPA, is the correlation between the observed phenotype and the predictions from a particular trained model fit on *n* samples (**Figure 1A**). In the simulated example, we train a ridge regression model on a sample of 500, yielding a true TPA of 0.41. TPA is dependent on the sample used to train the model and will fall anywhere less than or equal to MAPA. While MPA serves as a theoretical benchmark, MAPA quantifies the best predictive accuracy we could achieve for a given pipeline and is thus of most practical interest.

### Pearson’s correlation estimator is biased and has invalid confidence intervals

Pearson’s correlation estimator is the most common prediction accuracy estimator reported for ML brain-phenotype associations^10^. We evaluate the performance of the Pearson estimator and Fisher CI to capture MAPA using the simulated example above (**Figure 1B**).

We perform 5-fold cross-fitting as described above (also see **Methods**). The final Pearson estimate is the mean of the fold-specific estimates. Pearson’s estimator is downward biased for MAPA. While the estimator converges to MAPA in sample sizes above 2,500, the estimator performs poorly in samples reasonably accessible in most neuroimaging studies, which are typically less than 100^20^. The 95% Fisher CI^21^ fails to capture MAPA accurately, even for large sample sizes.

To assess the performance of Pearson’s estimator for TPA, we train a model outside of the simulated datasets on a set of 500 individuals, representing a pre-trained model. Within the simulations, we resample from the rest of the dataset not used to fit the model. Pearson’s estimator has small bias for this quantity (**Figure 1A**). The corresponding Fisher CI achieves the expected range of coverage given the number of simulations (**Figure 1B**). Critically, Pearson’s estimator does not represent the MAPA for the ML model class.

### Improved estimation of MAPA with semiparametric statistics

We provide an improved, semiparametric (also called double ML) one-step estimator for MAPA. A one-step estimator provides a first-order bias correction based on the efficient influence function of a parameter^22–27^. Our estimator adds a bias adjustment term to the empirical Pearson’s correlation estimate and applies a transformation to ensure the estimator respects the boundaries of the parameter space and improve small sample performance (**Methods**). The estimator is asymptotically normal with a straightforward CI procedure. We compare the performance of our estimator to the standard Pearson estimator and Fisher CI across a range of settings using simulated data. We developed several semiparametric estimators and selected one based on its finite sample performance in a separate dataset from the Adolescent Brain Cognitive Development (ABCD) study^28^ (**Figures S1, S3**).

To evaluate the one-step estimator, we first compare the performance of the estimators in plasmode simulations predicting age using the same regional GMVs from the DKT atlas as features in ridge and random forest models. The one-step estimator converges to MAPA faster than Pearson’s estimator for both ridge and random forest models across a range of MAPA values (**Figure 2A**, **Table S2**). Both estimators converge faster with the simpler model, ridge regression. For higher MAPA, the difference in convergence rate between ridge and random forest is less pronounced for the one-step estimator. For ridge regression, the expected value of the one-step estimator converges by approximately *n* = 5,000 for low MAPA and *n* = 3,000 for higher MAPA. In contrast, the expected value of Pearson’s estimator in ridge regression reaches MAPA by *n* = 20,000 for larger MAPA but is still biased at *n* = 20,000 for lower MAPA. For random forest, the one-step estimator reaches MAPA by *n* = 20,000 for higher MAPA, while Pearson’s estimator is still notably biased at *n* = 20,000 for all values of MAPA.

**Figure 2.**
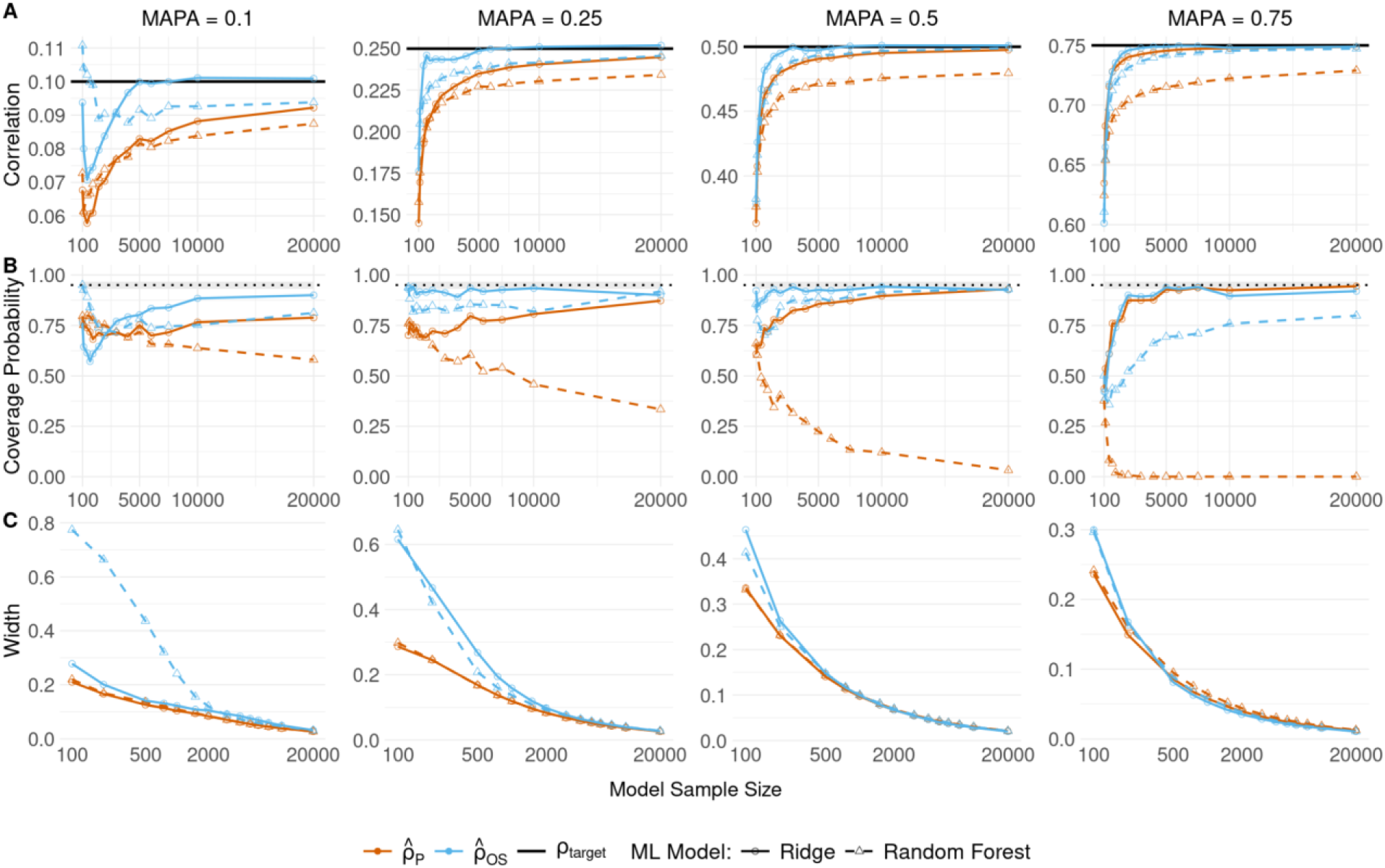
The semiparametric one-step estimator outperforms Pearson’s estimator across ML models in realistic plasmode simulations of anatomical (62 features) association with age. **A)** Mean estimates of the Pearson (orange) and one-step (blue) correlation across 500 simulations of ridge regression (solid line, circles) and random forest (dashed line, triangles) models for different values of MAPA, increasing from left to right in the columns. The one-step estimator is less biased and converges faster than the Pearson estimator. Convergence is faster for larger MAPA and simpler models (i.e., ridge). The one-step estimator has more stable performance across model types. **B)** 95% confidence interval coverage probability using the Fisher interval (for Pearson) or the one-step confidence interval. The gray shaded region is the 95% probability interval for the 500 simulations. The Pearson CI only reaches the nominal level with the simpler ridge model, and the CI coverage for the more complex random forest model goes to 0. **C)** Average confidence interval width for each method. The one-step CI is wider in small samples, but similar or narrower asymptotically than the Pearson CI.

We also evaluate the accuracy of 95% CI coverage probability and width (**Figure 2B-C**). Ninety-five percent CIs should capture the true parameter at least 95% of the time – of those CIs that do, the best has the smallest width. The one-step CI reaches nominal or near-nominal coverage in ridge regression across MAPA values, in some cases reaching the nominal level in sample sizes in samples of 1,000 or less (MAPA = 0.25 and 0.5). The Fisher CI for Pearson’s estimator has increasing coverage with increasing sample size and MAPA but only achieves similar performance to the one-step estimator for high MAPA (MAPA = 0.75). For random forest, the Fisher CI procedure fails, with coverage decreasing with increasing sample size and MAPA. The one-step CI has better coverage than Fisher’s CI for random forest, with coverage increasing with sample size. The one-step CIs are wider on average for small sample sizes but are asymptotically similar or narrower than the Fisher CI.

To generalize these findings to resting state functional connectivity (FC) data with higher-dimensional imaging features, we compare the performance of the estimators in similar plasmode simulations as above using data from 3161 individual participants predicting age with FC of varying resolution. We use random forest models with the Schaefer 300 atlas and consider both the 7-network (28 features) and 17-network (153 features) mappings^29,30^. The one-step estimator shows lower bias than Pearson’s estimator and both estimators show faster convergence for the model using the lower-dimensional features (**Figure 3A**). CI coverage is closer to the nominal level for the one-step estimator and goes to zero with increasing sample size for Fisher’s CI (**Figure 3B**). These findings are consistent in analogous models using ridge regression (**Figure S4**) and demonstrate superiority of the one-step estimator. We perform the same simulations with psychiatric phenotypes and find consistent results (**Figure S5**).

**Figure 3.**
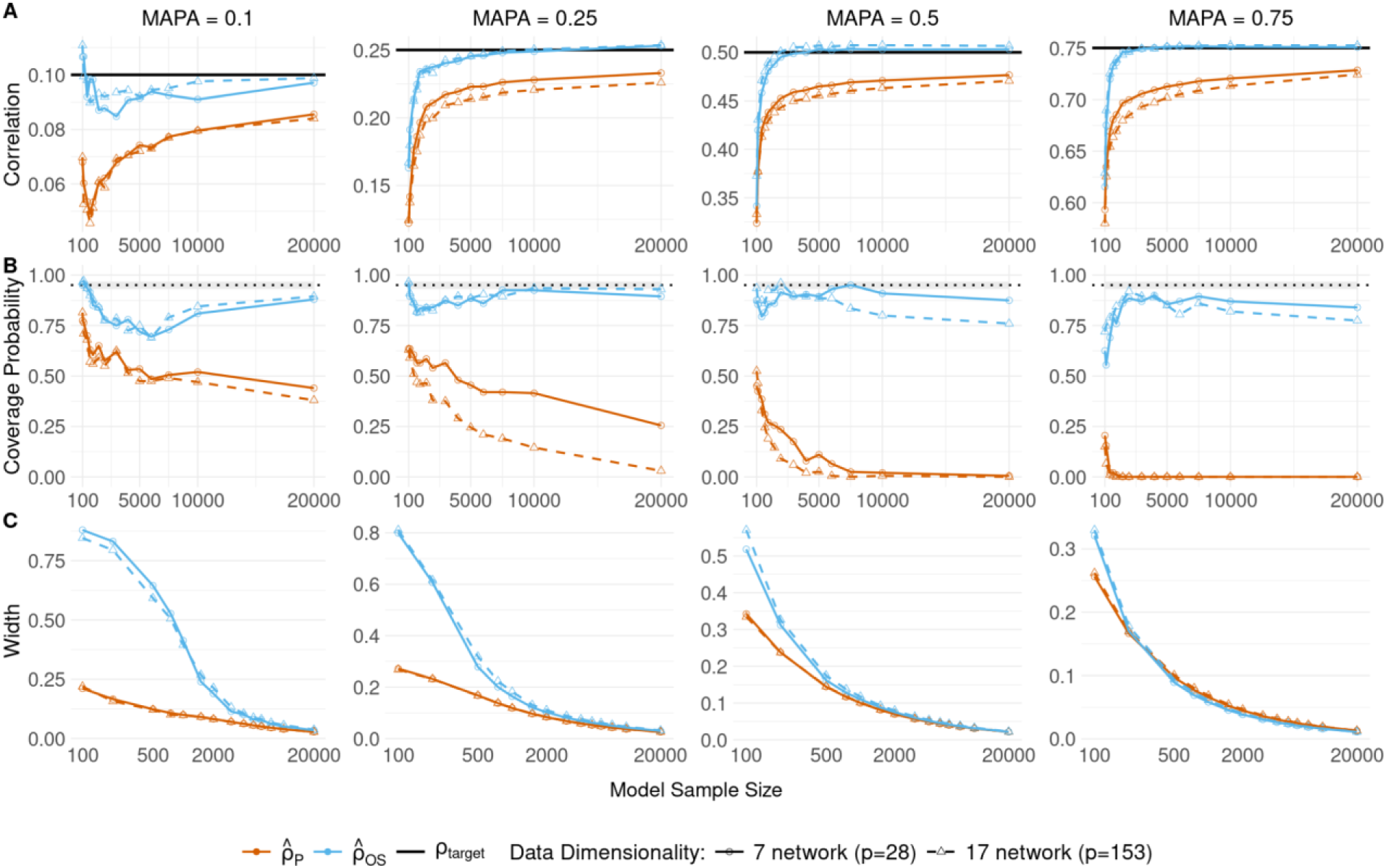
The one-step estimator outperforms Pearson’s estimator across data dimensionalities in plasmode simulations of functional connectivity with age. **A)** Mean estimates of the Pearson (orange) and one-step (blue) correlation for correctly specified and tuned random forest models using either 7 network FC data (solid line, circles; number of predictors = 28) or 17 network FC data (dashed line, triangles; number of predictors = 153) for different values of MAPA, increasing from left to right in the columns. The one-step estimator is less biased and converges faster than the Pearson estimator. Convergence is faster for stronger MAPA and less complex models (i.e., 7 networks). **B)** 95% confidence interval coverage probability using the Fisher interval (for Pearson) or the one-step confidence interval. The gray shaded region is the expected range of coverage for 500 simulations. The one-step CI outperforms the Pearson, reaching or near the nominal level asymptotically. **C)** Average confidence interval width for each method. The one-step CI is wider in small samples, but similar or narrower asymptotically than the Pearson CI.

### ML associations with age and psychopathology

We apply the one-step estimator to study brain-phenotype associations using neuroimaging data to predict age, P-factor score, externalizing score, and internalizing score in the full RBC dataset (*n* = 3074; **Table 1**; **Methods**). We compare ridge regression and random forest models using combinations of functional and structural neuroimaging features to a baseline model that includes nuisance features: age (for the psychiatric phenotypes), sex, Euler number, and mean frame displacement. The structural models include 62 regional GMV features from the DKT atlas and the baseline covariates. The functional models include 153 FC features from the 17-network Schaefer atlas and the baseline covariates.

For each model, we perform repeated 5-fold cross-fitting with nested model tuning, repeating the analyses across 250 sample splits to ensure stability and replicability of the point estimates and CIs (**Methods**). Final estimates and CIs for both estimators are the median value across all sample splits^25^. Structural and functional data are highly predictive of age and weakly predictive of psychopathologic phenotypes (**Figure 4**). The one-step estimates are larger than the Pearson’s estimates, with larger and positively skewed CIs, reflecting the true uncertainty of the ML estimates. Compared to ridge regression, the random forest models show improvement in prediction accuracy of varying magnitude depending on the phenotype.

**Figure 4.**
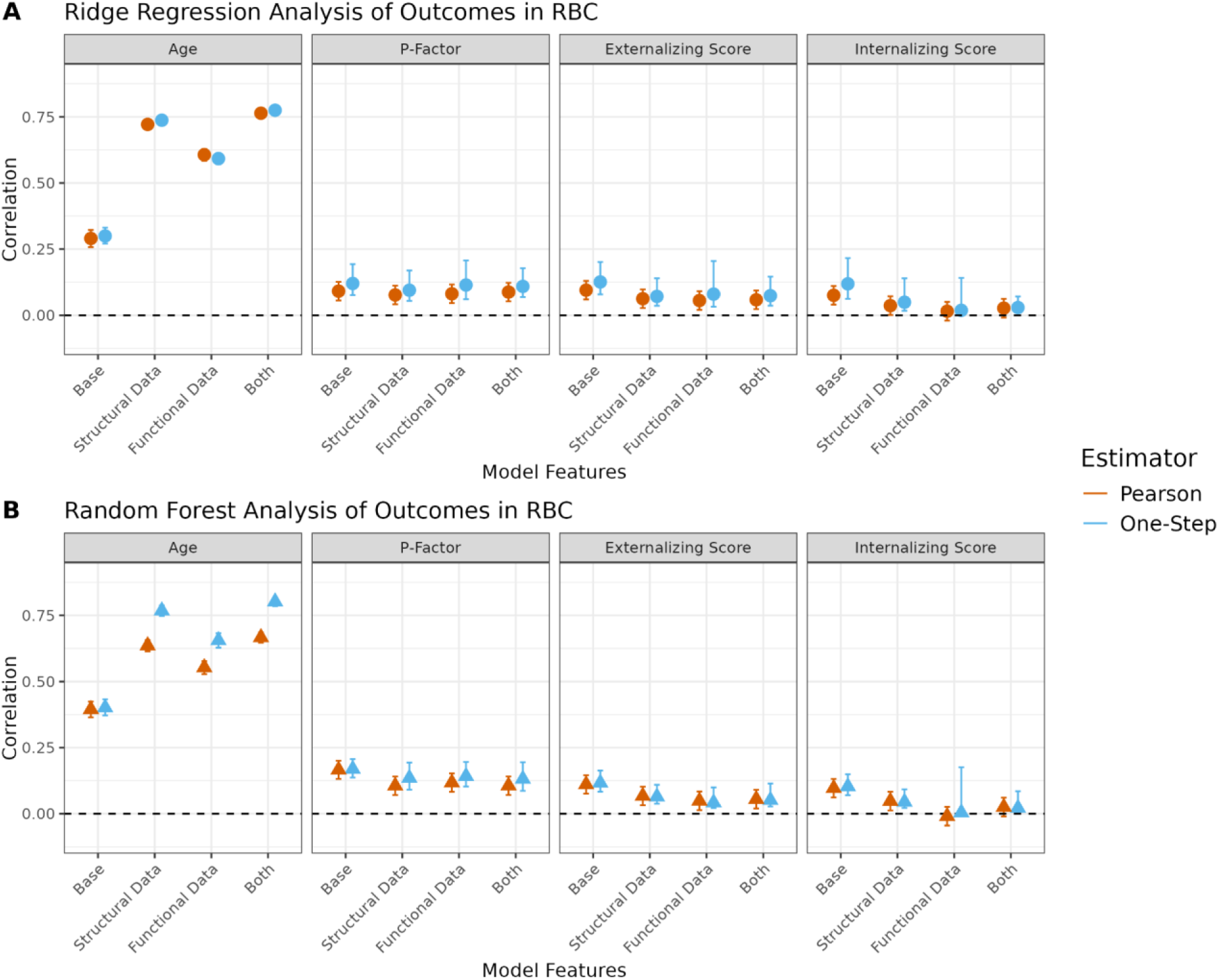
Functional and structural imaging data are highly predictive of age but do not improve prediction accuracy of psychopathologic phenotypes. Median estimates across 250 sample splits of the Pearson and one-step correlation for (**A**) random forest models and (**B**) ridge regression using no imaging data (“Base”), structural data, functional data, or both as model features predicting four outcome phenotypes, shown with 95% confidence intervals. For most associations, the one-step estimator has a slightly larger estimate and larger, positively skewed confidence interval than Pearson’s correlation. Random forest offers consistent, but varying improvement in prediction accuracy across all outcomes compared to ridge regression.

Unlike the Pearson estimator, the one-step estimator allows us to statistically compare the difference between competing models (**Figure 5**; **Methods**). Functional and structural data and their combination improve prediction performance over baseline models for age, with the structural and combination data performing better than functional data. For the psychopathology phenotypes, inclusion of imaging data has equal or lower prediction accuracy. We also compare models using an effect size ratio derived from the one-step approach and reach similar conclusions (**Figure S6**).

**Figure 5.**
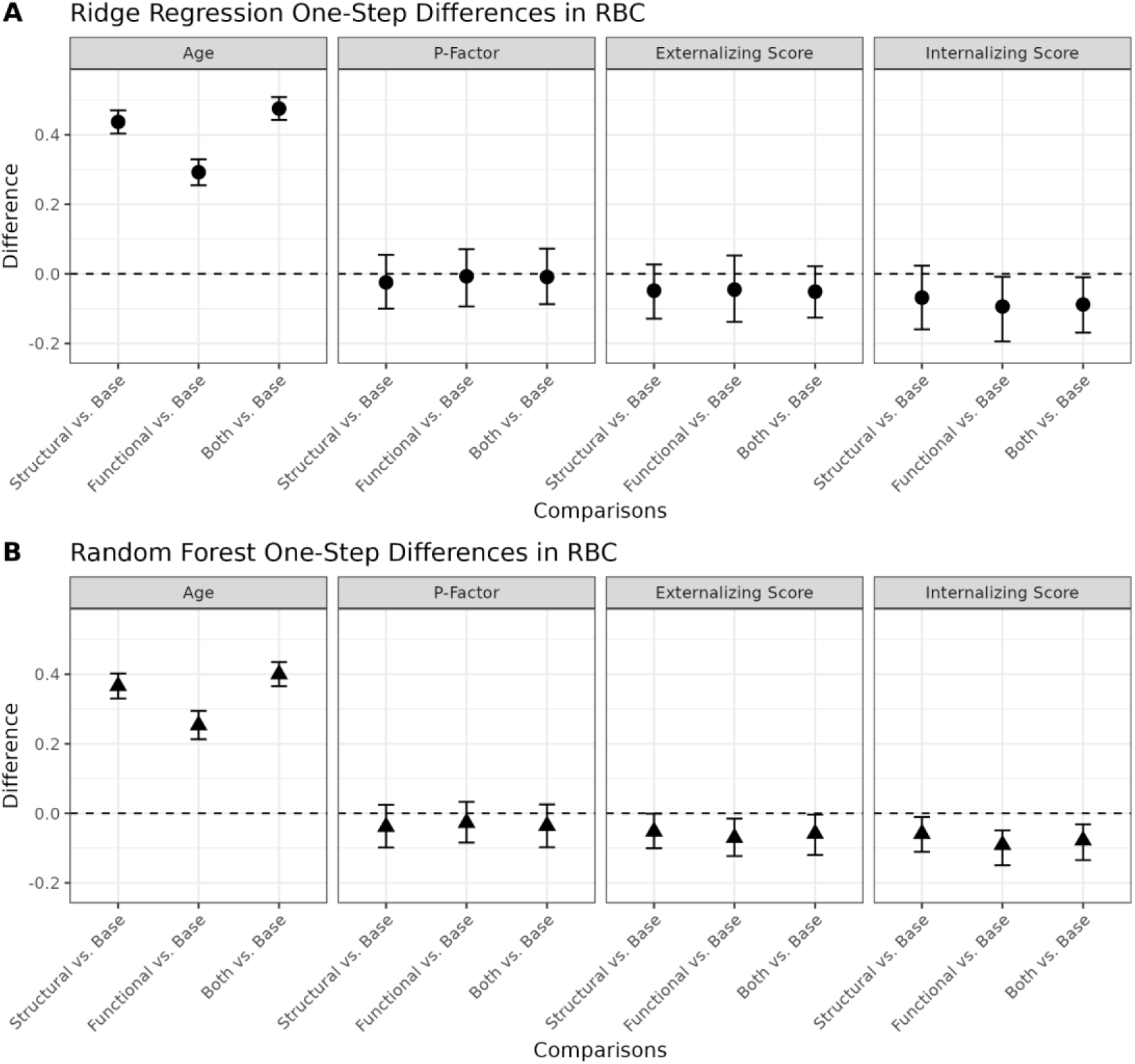
Inclusion of imaging data increases prediction accuracy for age. Median estimate across 250 sample splits comparing prediction accuracy between structural, functional, or both versus no imaging data (“Base”) for the one-step estimator using (**A**) random forest and (**B**) ridge regression for four outcome phenotypes. Random forest models with structural and a combination of functional and structural data perform best against random forest models with no imaging data when predicting age. The inclusion of imaging data in random forest models that predict psychopathologic outcomes results in decreased prediction accuracy. Ridge regression models with structural and a combination of functional and structural data perform best against ridge regression models with no imaging data when predicting age. The inclusion of imaging data in ridge regression models that predict psychopathologic outcomes results in decreased prediction accuracy.

To compare the empirical results of the one-step estimator to Pearson’s correlation in a typical NIH R01-scale sample, we repeat the analysis in a subset of the RBC data from one study site (HBN Staten Island, *n* = 208) and obtain similar findings (**Table S3**, **Figures S7, S8**). There are larger CIs due to higher uncertainty and boundary-related performance of the one-step estimator.

## Discussion

Explicit articulation of the goal of estimating predictive accuracy is paramount for replicable brain-phenotype associations. Studies that predict continuous phenotypes often report an estimate for the correlation of the trained model’s predictions with the observed phenotype, but the target of estimation is unclear, obfuscating comparisons between prediction accuracies. We highlight the distinction between the MAPA and the prediction accuracy of a single trained model, TPA, emphasizing that the latter is dataset and sample size dependent. Likewise, we show that the standard Pearson correlation estimator is tied to the specific data (and sample size) used to train the model. It systematically underestimates the target correlation, with worse performance for weaker true effect sizes, more flexible ML methods, and higher-dimensional input data. Additionally, standard confidence interval procedures fail to capture the target correlation. We propose a semiparametric one-step estimator and confidence interval for MAPA and show that it performs considerably better across a range of data types, ML algorithms, and data dimensionalities. In comparison to standard practice, broad use of this estimator would, for the first time, allow accurate measurement and comparison of MAPA in efforts to link brain and behavior using ML tools.

The one-step estimator allows comparison of MAPA for two ML models. We used this to compare models with regional and network imaging features to a base model including only image quality features and demographic variables. Critically, these results indicate imaging measures do not increase MAPA for these aggregate psychopathology phenotypes above what can be ascertained from demographic variables. Our results were consistent with prior literature that did not control for covariates^31^ and suggest these phenotypes may be too heterogeneous to be consistently associated with structural and functional MRI features^32,33^. More specific phenotypes or alternative imaging measures that are less processed using more complex deep learning models^7^ may yield higher MAPA.

Replicability is an ongoing concern for brain-phenotype associations, where effect size estimates are often unstable or biased^1,34,35^. Our findings demonstrate that, for ML analyses, standard methods underestimate the true effect and are highly dependent on training sample size. This undermines replicability and complicates comparisons across studies^36^. By providing less biased estimates of MAPA for a given brain-phenotype association and ML model, the one-step estimator yields more realistic expectations for the strength of underlying effects. The MAPA for an ML class is a lower bound for the true predictive potential for a given brain-phenotype association^9^. Modeling choices may result in a model class that is better able to approximate the true underlying relationship. Thus, thorough reporting of ML methods is critical to parse heterogeneity across studies.

The one-step estimator accommodates inference for MAPA in feasible large-scale neuroimaging studies. For ridge regression models, we observe low bias in sample sizes ranging from 1,000 to 20,000, although our method still outperforms the standard method in smaller samples. The sample size needed for unbiased estimation depends on the magnitude of the true effect and the complexity of the model. Current guidelines often caution against training a predictive neuroimaging model with less than 200-500 participants^8,37,38^, but our results suggest MAPA will likely still be underestimated with these sample sizes. The one-step estimator substantially reduces this “small-sample” bias, offering greater accuracy across studies.

Quantifying uncertainty around predictive accuracy estimates is crucial to better understand the potential magnitude of brain-phenotype associations and the stability of the estimator. Confidence intervals for correlation are not often reported for ML brain-phenotype associations^10^, and our results show that the standard procedure does not have proper coverage. The one-step estimator improves the confidence intervals but can still have under nominal coverage. When the true effect size is close to zero, our method can produce very wide CIs, sometimes spanning 0 to 1 (**Figure S7**) due to the ML model producing near-constant predictions. In this scenario, a permutation test can complement the interpretation of the CI. Further research could investigate improvements for small samples, perhaps by considering higher-order influence functions^39,40^ (**Methods**).

Model tuning is critical to ensure ML models have convergence rates required for valid inference using semiparametric methods (**Methods**)^26^. These methods relax assumptions on the convergence rate of the ML model compared to Pearson’s correlation, but they do not remove complexity constraints entirely. With proper hyperparameter tuning, many highly flexible models can achieve the needed convergence rate (**Table S2**). We recommend a second level of cross-validation, nested within the cross-fitting procedure, to perform model tuning. It is advisable to repeat the cross-fitting procedure many times (e.g., 250)^25^ over different splits of the sample to improve reproducibility and replicability for estimating MAPA.

Our work establishes the importance of using modern statistical theory to quantify accuracy in ML brain-phenotype associations. When investigating potential models for relationships between brain data and individual traits, it is important to quantify an estimate of the MAPA of the model to evaluate the utility of the imaging modality for predicting the phenotype. Our method, which accounts for the randomness of the training data and adjusts for sample-size-dependent bias, allows researchers to more accurately quantify achievable predictive accuracy using modern black-box ML techniques. This opens the door to benchmarking predictive potential across analytic choices, accelerating replicable discovery in ML brain–phenotype research.

## Methods

### Data and preprocessing

We analyzed data from the Reproducible Brain Charts (RBC) open data resource and used these data to conduct realistic plasmode simulations^15,41^. RBC aggregated data from five neuroimaging studies, including the Developmental Chinese Color Nest Project (CCNP^42^; n = 195), the Philadelphia Neurodevelopmental Cohort (PNC^43^; n = 1,601), the Nathan Kline Institute-Rockland Sample (NKI^44^; n = 1,329), the Healthy Brain Network (HBN^45^; n = 2,611), and the Brazilian High Risk Cohort (BHRC^46^; n = 610). RBC data was deidentified and collected with IRB approval from the institutions associated with each study^41^.

RBC performed rigorous quality assurance and applied the same preprocessing pipeline to all data to maximize comparability. Details of image acquisition and preprocessing are available in prior work^41^. Briefly, preprocessing was performed using the FAIRly-big strategy^47^. Structural MRI scans were corrected for intensity non-uniformity and skull-stripped using ANTs in sMRIPrep and subsequently reconstructed with FreeSurfer. Functional data was processed via the Configurable Pipeline for the Analysis of Connectomes (C-PAC)^48^, with a custom configuration available online (https://reprobrainchart.github.io/docs/imaging/image_processing/). Processed data are publicly available for a variety of outputs and brain atlases (https://reprobrainchart.github.io/). The RBC includes harmonized psychiatric phenotypes from a bifactor model from the Child Behavior Checklist (CBCL), used by CCNP, NKI, HBN, and BHRC; and GOASSESS, used by PNC^49–52^.

We included only baseline data for multi-session studies (NKI and BHRC). For structural analyses, we downloaded tabulated regional gray matter volume from the Desikan-Killiany-Tourville atlas^16^. For functional data, we downloaded the functional connectivity matrices parcellated using the Schaefer 300 atlas with 36-parameter preprocessing regression corresponding to the earliest run passing QC^29,30^. To calculate network-to-network connectivity, we took the mean of the correlations for regions within and between networks. We only downloaded data passing the strictest quality threshold (complete pass)^41^. For plasmode simulations, we included all downloaded scans (**Simulation methods**). For the phenotype association analyses, we used only the subset of participants that had both structural and functional imaging data that passed quality control.

To account for differences in data acquisition across sites, we applied CovBat to harmonize the means and covariances of imaging data while preserving relationships with model covariates using the default parameters, with the number of principal components harmonized set to explain 95% of the variance^53^. We considered each study site as a batch (BHRC has 2 sites, and HBN has 4 sites; the other studies are single-site) and included age, sex, P-Factor, internalizing score, externalizing score, and attention score as covariates. We harmonized structural and functional imaging separately.

### Statistical theory

#### Notation

Let *X* denote a set of features (e.g., region-region edges from FC network) and *Y* the phenotype of interest (e.g. age), collected on *n* individuals. We denote the true population conditional mean, without making complexity assumptions, with ξ. In practice, we restrict to a class of models *M* and denote *μ* as the best approximation to ξ within this class.

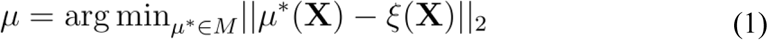

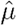 denotes the trained predictive model for a specific training set.

#### Parameter Definitions

For continuous phenotypes, the correlation between predicted and observed values is often taken as a metric of predictive accuracy^10^. Using the notation above, we define three different, but related, correlation parameters:

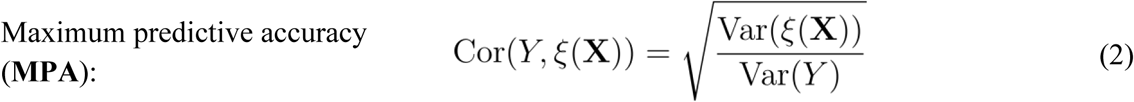

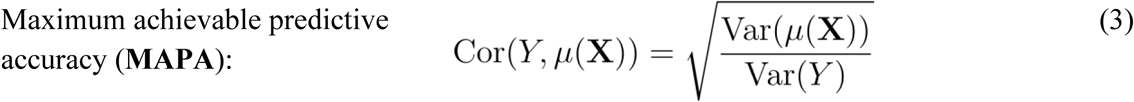

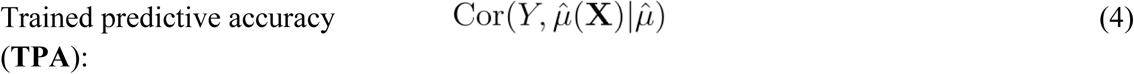

We define MPA as the correlation between the phenotype and the true mean model, while the MAPA for a model class is the correlation between the observed phenotype and the predictions from the closest model within the restricted class. Because we express MPA and MAPA as the root of the ratio of the variance of the conditional mean (true or restricted, respectively) to the variance of the phenotype, the parameter space is [0,1]. Trained predictive accuracy (TPA) denotes the correlation between the observed phenotype and the predicted values from a particular fitted model, assuming the predicted model to be fixed and known. TPA is dependent on the characteristics, including sample size, of the training set. In general, MPA ≥MAPA ≥ TPA.

#### Pearson estimator and Fisher confidence interval

Pearson’s correlation is a typical plug-in estimator for any of these three parameters,

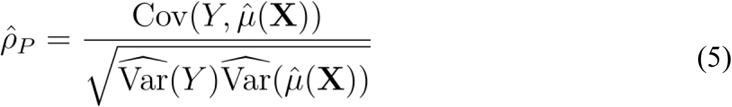

Training/testing data split, cross-validation, or cross-fitting is used to estimate MAPA (**Statistical analyses:** *Cross-fitting*). Given the simplification of the parameter for prediction (Equation 3), we truncated our estimator of MAPA at 0 to obtain an estimator with lower mean square error. To estimate TPA, a pretrained model can be used to generate predictions for the full dataset and compute Pearson’s correlation. We used Fisher’s transformation^21,54^ to compute a CI for Pearson’s correlation.

#### One-step estimator and confidence interval

Estimators of MAPA depend inherently on the randomness and complexity of estimating the *μ* function. Using flexible ML methods and high-dimensional neuroimaging methods, we cannot assume that 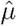 will converge to *μ* at the parametric 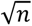-rate generally required for standard central limit theorems to apply^26,27^. Additionally, the plug-in Pearson estimator may suffer from bias due to slow convergence of the ML model^25,26^. These problems motivate development of an improved estimator for MAPA.

We adopted a semiparametric framework to develop a one-step estimator for MAPA. One-step estimators correct a plug-in estimate by adding a term based on the empirical mean of the efficient influence function of the target quantity^22–27^. Semiparametric theory identifies conditions when one-step estimators are asymptotically normal and the variance can be validly estimated as the empirical variance of the empirical influence function (**Supplement Section S4**). We derived the influence function for a transformed version of MAPA to ensure the estimator respects the [0,1] bounds. Our adjusted estimator is

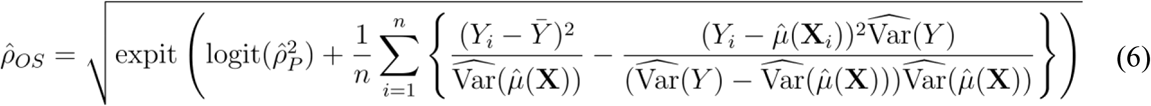

We computed CIs for the estimator as

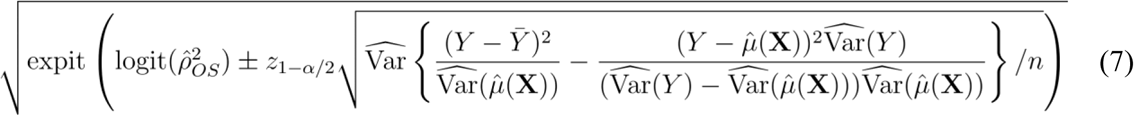

We investigated many transformations and variations on the estimator formula, with the above approach yielding the best small sample performance (**Supplement Section 2.3**; **Table S1**; **Figures S1, S3**). We confirmed the mean estimated standard error is close to the true standard error of the one-step estimator asymptotically across simulations (**Figure S2**).

Leveraging the asymptotic normality of the one-step estimator, we employed the multivariate Delta Method^27^ to construct estimates and CIs for the difference in MAPA for different model classes (**Supplementary Section S7**).

### Statistical analyses

#### Phenotype associations in the RBC

To evaluate empirical differences between methods, we modeled age and three of the psychiatric phenotypes available in the RBC (p-factor, internalizing score, and externalizing score). For each phenotype, we fit four models: (1) baseline covariates only (age (for non-age phenotypes), sex, mean frame displacement, and Euler number), (2) structural data plus baseline covariates (66 features), (3) functional data plus baseline covariates (157 features), and (4) structural plus functional and baseline covariates (219 features). We included the imaging quality control measures in the baseline model to evaluate only the incremental value of the imaging features. These models were fit to the full harmonized dataset and a subset of the HBN data from the Staten Island site to evaluate the behavior of the methods in sample sizes of typical neuroimaging studies (**Table S3**; **Data and preprocessing**)^20^.

#### ML model training and inference

We evaluated the performance of our estimator using ridge regression and random forest. All analyses were performed in R version 4.4.1 using the glmnet, ranger, and tuneRanger packages^55–58^. Overfitting with ML models was mitigated using sample-splitting and cross-fitting^25,26^. For cross-fitting, we split the dataset into five folds and used the data from all but one fold to train the model – computing predictions, influence function estimates, and correlation estimates on the left-out fold – and then repeating the procedure for all folds. This ensured data used to train the model is independent of the data used to compute the influence function, so we avoided “double-dipping.” The final point estimate was taken as the mean of the five fold-specific estimates. The variance of the final estimate was computed by pooling the influence function estimates from all folds in a single vector and calculating the empirical variance. For estimating TPA, we split the resampled test data into five folds to maintain sample size consistency with the cross-fitting procedure described above, but we did not retrain the model. In each fold, the predictions and correlation estimates are computed from the given pretrained model.

Parameter tuning helps ensure that the ML models satisfy assumptions needed for approximate normality using the one-step estimator (**Supplementary Section S4**). We performed nested model tuning within each training dataset. For ridge regression, we performed 10-fold cross-validation to select the penalty parameter to minimize squared error loss from a grid, with finer steps approaching 0 ({10, 9, …, 1, 0.9, …, 0.1, 0.09, …, 0.01, 0.009, …, 0.001, …, 0.0009, …, 0.0001, …, 10^−7, 0}). For random forest models, we tuned the minimal node split size, the fraction of observations to sample, and the number of variables to split on via model-based optimization to minimize out-of-bag mean squared error. We used 25 tuning iterations after 10 warm-up iterations. All random forest models used 250 trees. For all models, if the tuned model gave constant predictions, we reverted to the nearest hyperparameter settings that did not yield constant predictions in order to compute the correlation estimates.

Because the cross-fit folds are dependent on randomly splitting the data, we performed multiple sample splits to ensure stability and reproducibility of the point estimates and CIs^25^. For simulations, because the results of the analyses were analyzed over 500 simulations, and due to computational complexity, we did a single sample split for cross-fitting. In the RBC analysis, we performed 250 sample splits and reported the final estimator and CIs as the median of the cross-fitted values across all splits.

### Simulation methods

We used the RBC data to perform realistic plasmode simulations. For each simulation setting, we split the data into a build dataset (20%) and an evaluation dataset. Using the build data, we fit an ML model using the real observed features (e.g., structural or functional data) to predict the phenotype (**Statistical analyses**: *Phenotype associations in the RBC*). We then applied this model to the evaluation dataset and considered the predicted values to be the “true” conditional mean of the phenotype. In each simulation replicate, we resampled the feature data, and corresponding conditional mean, from the evaluation dataset with replacement. Then, for each resampled participant, we generated the observed phenotype as the conditional mean plus a randomly sampled error, drawn from a uniform distribution with variance controlled to fix the value of MPA (**Supplementary Section S8**). For each resampled dataset, we performed 5-fold cross-fitting (with nested tuning) to compute estimates for Pearson’s estimator and our one-step estimator, with corresponding CIs (**Statistical analyses**: *Model training and inference*). We used resampled datasets of 14 sample sizes: *n* = 100, 200, 500, 750, 1000, 1500, 2000, 3000, 4000, 5000, 6000, 7500, 10000, 20000.

For **Figure 1**, to illustrate the theoretical differences between parameters, we used a random forest model to generate the “true” conditional mean (corresponding to MPA) and fit ridge regression models to the simulated data (corresponding to MAPA, with the parameter value computed via the ordinary least squares fit of the true conditional mean in the full evaluation dataset, representing the limit of ridge regression). For TPA, we trained one ridge regression model in 500 samples from the evaluation dataset. Then, in each resample of the remaining data, we computed the correlation between the fixed model’s predictions and the true conditional mean, adjusted for variability in the phenotype (**Supplementary Section S9**). We defined the ground truth TPA as the average over 500 simulations at the largest simulated sample size, *n* = 20,000. For subsequent simulation analyses, we assumed MPA = MAPA, meaning that the model in the build data was of the same class as the model we trained in the evaluation data (e.g., a ridge regression build model was used for the ridge regression simulations).

## Data Availability

The data we used in this work can be downloaded from GitHub (https://github.com/orgs/ReproBrainChart/repositories), following instructions on RBC’s website (https://reprobrainchart.github.io/docs/get_data).

For supplementary analyses, we used data obtained from the Adolescent Brain Cognitive Development^SM^ (ABCD) Study (https://abcdstudy.org), held in the NIMH Data Archive (NDA). This is a multisite, longitudinal study designed to recruit more than 10,000 children age 9-10 and follow them over 10 years into early adulthood. The ABCD Study® is supported by the National Institutes of Health and additional federal partners under award numbers U01DA041048, U01DA050989, U01DA051016, U01DA041022, U01DA051018, U01DA051037, U01DA050987, U01DA041174, U01DA041106, U01DA041117, U01DA041028, U01DA041134, U01DA050988, U01DA051039, U01DA041156, U01DA041025, U01DA041120, U01DA051038, U01DA041148, U01DA041093, U01DA041089, U24DA041123, U24DA041147. A full list of supporters is available at https://abcdstudy.org/federal-partners.html. A listing of participating sites and a complete listing of the study investigators can be found at https://abcdstudy.org/consortium_members/. ABCD consortium investigators designed and implemented the study and/or provided data but did not necessarily participate in the analysis or writing of this report. This manuscript reflects the views of the authors and may not reflect the opinions or views of the NIH or ABCD consortium investigators. The ABCD data repository grows and changes over time. This study used ABCD Data Release 5.1 available at (http://dx.doi.org/10.15154/z563-zd24).

## Code Availability

The code for all simulations and analyses can be found at https://github.com/statimagcoll/Semiparametric_Correlation. Additionally, the R package **semicor** (https://github.com/statimagcoll/semicor) has functions to compute the one-step correlation estimator and confidence interval using ML model predictions.

## Acknowledgements

This research was supported by NIH grants R01MH133843 (to AA-B) and R01MH123563 (to SNV).

## Author Contributions

M.T.J., E.K., and S.V. conceptualized the work. M.T.J., J.L., E.K., and S.V. developed the methodology. M.T.J. and S.V. conducted the simulations. M.T.J, I.G., and S.V. conducted the real data analysis. M.T.J and I.G. wrote the R package. M.T.J. and K.K. performed data processing, with input from A.C. X.Z. performed code validation. M.T.J., I.G., A.C., J.S., A.A-B., and S.V. interpreted the results. M.T.J., I.G., and S.V. drafted the manuscript. All authors revised and edited the manuscript.

## Competing Interests

J.S. and A.A-B. hold equity in and J.S. is a director of Centile Bioscience.

## Notes

https://github.com/statimagcoll/Semiparametric_Correlation

https://github.com/statimagcoll/semicor

