## Supplement for "Improved Estimation of Correlation Accuracy for Machine Learning Brain-Phenotype Associations"

November 26, 2025

#### Contents

**S1 Parameter definitions** 2

**S2 Influence functions for MAPA** 3
S2.1 Full influence function 3
S2.2 Mixed Influence Function 4
S2.3 Parameter transformations 4
S2.3.1 Squared logit transformation 4
S2.3.2 Logit transformation 4
S2.3.3 Fisher transformation 5

**S3 Constructing one-step estimators** 5
S3.1 Evaluating parameter transformations 5

**S4 Requirements for asymptotic normality** 6

**S5 Methods to combine estimators across folds** 7

**S6 Additional simulation results** 8
S6.1 Evaluation of dimensionality with functional connectivity data 8
S6.2 Evaluation of psychiatric prediction with structural data 9

**S7 Comparing models via the Delta Method** 10
S7.1 Comparing the difference in correlations 11
S7.2 Comparing the effect size ratio 11

**S8 Simulation error distribution** 12

**S9 Computing ground truth TPA** 13

**S10 Single-sample evaluation in HBN Staten Island site** **13**

**S11 One-step estimator formulas** **16**

### S1 Parameter definitions

As in the paper, let $X$ denote the imaging features and $Y$ the continuous phenotype of interest, collected on $n$ individuals. Denote $\xi(\mathbf{X}) = E[Y|\mathbf{X}]$, the true conditional mean of the phenotype given the imaging data. We define maximum achievable predictive accuracy (MAPA) as the maximum achievable accuracy within a class of models, $M$, that we will fit to the data. Denote $\mu$ as the best approximation to $\xi$ within the class $M$, defined as the L2 projection of $\xi$ onto the space of models spanned by $M$,

$$\mu(\mathbf{X}) = \arg\min_{\mu^* \in M} ||\mu^*(\mathbf{X}) - \xi(\mathbf{X})||_2$$

The correlation (i.e., maximum predictive accuracy) of the phenotype with the true conditional mean can be simplified to

$$\begin{aligned}
\text{MPA} = \rho_0 &= \text{Cor}(Y, \xi(\mathbf{X})) \\
&= \frac{\text{Cov}(Y, \xi(\mathbf{X}))}{\sqrt{\text{Var}(Y)\text{Var}(\xi(\mathbf{X}))}} \\
&= \frac{\text{E}[\xi(\mathbf{X})^2] - E[Y]^2}{\sqrt{\text{Var}(Y)\text{Var}(\xi(\mathbf{X}))}} \\
&= \sqrt{\frac{\text{Var}(\xi(\mathbf{X}))}{\text{Var}(Y)}}
\end{aligned}$$

where the simplification follows from the law of iterated expectations.

Replacing $\xi$ with $\mu$ above yields the correlation of the phenotype with the best approximation to the conditional mean within the class under consideration (i.e., the maximum achievable predictive accuracy),

$$\text{MAPA} = \rho_1 = \text{Cor}(Y, \mu(\mathbf{X})) \quad (1)$$

$$= \sqrt{\frac{\text{Var}(\mu(\mathbf{X}))}{\text{Var}(Y)}}. \quad (2)$$

The second line, (2) follows by the orthogonality of $\xi(X) - \mu(X)$ due to $\mu(X)$ being an $L^2$ projection onto $M$. We can also quantify the relationship between MPA and MAPA:

$$\begin{aligned}
\rho_1 &= \text{Cor}(Y, \mu(\mathbf{X})) \\
&= \text{Cor}(Y, \xi(\mathbf{X})) \times \text{Cor}(\xi(\mathbf{X}), \mu(\mathbf{X})) \\
&= \rho_0 \times \text{Cor}(\xi(\mathbf{X}), \mu(\mathbf{X})).
\end{aligned}$$

The MAPA within a class of models is equal to the product of the correlation between the phenotype and the true conditional mean and the correlation between the true conditional mean and its projection.

Denote a particular trained predictive model trained on $(\mathbf{X}_i, Y_i)_{i=1}^n$ as $\hat{\mu}$. The correlation between the phenotype and the predictions from this trained model is the trained predictive accuracy,

$$\text{TPA} = \text{Cor}(Y, \hat{\mu}(\mathbf{X})|\hat{\mu}).$$

#### S2 Influence functions for MAPA

The key to deriving semiparametric one-step estimators is to find the influence function (IF) for the parameter of interest[1–6]. Here, we derive the IFs for MAPA and later consider transformations. Starting from (1) or (2), yields the same IF.

We can derive IFs for pathwise differentiable parameters, such as those above, with rules analogous to derivative rules in calculus[5] and by breaking down the full IFs into functions of IFs of simpler parameters. Define $\mathbb{IF}$ to indicate the influence function operator. We use the following IF formulas[5]:

- $\mathbb{IF}(\mathrm{E}[X]) = X - \mathrm{E}[X]$
- $\mathbb{IF}(\mathrm{Var}(X)) = (X - \mathrm{E}[X])^2 - \mathrm{Var}(X)$
- $\mathbb{IF}(\mathrm{E}[Y|X]) = \frac{\mathbf{1}(X=x)}{p(X=x)}\{Y - \mathrm{E}[Y|X]\}$
- $p(X = x) = \mathbf{1}(X = x) - p(X = x)$

##### S2.1 Full influence function

To derive the IF for MAPA, we start with the chain rule:

$$\begin{aligned}
\mathbb{IF}(\rho_1) &= \mathbb{IF}\left(\frac{\sqrt{\mathrm{Var}(\mu(\mathbf{X}))}}{\sqrt{\mathrm{Var}(Y)}}\right) \\
&= \frac{1}{2}\left(\frac{\mathrm{Var}(\mu(\mathbf{X}))}{\mathrm{Var}(Y)}\right)^{-\frac{1}{2}} \mathbb{IF}\left(\frac{\mathrm{Var}(\mu(\mathbf{X}))}{\mathrm{Var}(Y)}\right) \\
&= \frac{1}{2}\left(\frac{\mathrm{Var}(Y)}{\mathrm{Var}(\mu(\mathbf{X}))}\right)^{\frac{1}{2}} \left(\frac{\mathbb{IF}(\mathrm{Var}(\mu(\mathbf{X})))\mathrm{Var}(Y) - \mathbb{IF}(\mathrm{Var}(Y))\mathrm{Var}(\mu(\mathbf{X}))}{\mathrm{Var}(Y)^2}\right) \qquad (3)
\end{aligned}$$

Using the building blocks above, we derive the influence function for $\mathrm{Var}(\mu(\mathbf{X}))$:

$$\begin{aligned}
\mathbb{IF}(\mathrm{Var}(\mu(\mathbf{X}))) &= \mathbb{IF}(\mathrm{E}[\mu(\mathbf{X})^2] - \mathrm{E}[\mu(\mathbf{X})]^2) \\
&= \mathbb{IF}(\mathrm{E}[\mu(\mathbf{X})^2]) - \mathbb{IF}(\mathrm{E}[Y]^2) \\
&= \mathbb{IF}\left(\sum_{\mathbf{x}} \mu(\mathbf{x})^2 p(\mathbf{x})\right) - 2\mathrm{E}[Y]\mathbb{IF}(\mathrm{E}[Y]) \\
&= \sum_{\mathbf{x}} \mathbb{IF}(\mu(\mathbf{x})^2 p(\mathbf{x})) - 2\mathrm{E}[Y]\{Y - \mathrm{E}[Y]\} \\
&= \sum_{\mathbf{x}} \left\{\mathbb{IF}(\mu(\mathbf{x})^2)p(\mathbf{x}) + \mu(\mathbf{x})^2\mathbb{IF}(p(\mathbf{x}))\right\} - 2\mathrm{E}[Y]\{Y - \mathrm{E}[Y]\} \\
&= \sum_{\mathbf{x}} \left\{2\mu(\mathbf{x})\mathbb{IF}(\mu(\mathbf{x}))p(\mathbf{x}) + \mu(\mathbf{x})^2\left(\mathbf{1}(\mathbf{X} = \mathbf{x}) - p(\mathbf{X} = \mathbf{x})\right)\right\} - 2\mathrm{E}[Y]\{Y - \mathrm{E}[Y]\} \\
&= \sum_{\mathbf{x}} \left\{2\mu(\mathbf{x})\left(\frac{\mathbf{1}(\mathbf{X} = \mathbf{x})}{p(\mathbf{X} = \mathbf{x})}\{Y - \mathrm{E}[Y|\mathbf{X}]\}\right)p(\mathbf{x}) + \mu(\mathbf{x})^2(\mathbf{1}(\mathbf{X} = \mathbf{x}) - p(\mathbf{X} = \mathbf{x}))\right\} \\
&\quad - 2\mathrm{E}[Y]\{Y - \mathrm{E}[Y]\} \\
&= \sum_{\mathbf{x}} \{2\mu(\mathbf{x})\mathbf{1}(\mathbf{X} = \mathbf{x})Y - 2\mu(\mathbf{x})\mathbf{1}(\mathbf{X} = \mathbf{x})\mu(\mathbf{x}) + \mu(\mathbf{x})^2\mathbf{1}(\mathbf{X} = \mathbf{x}) - \mu(\mathbf{x})^2 p(\mathbf{X} = \mathbf{x})\} \\
&\quad - 2\mathrm{E}[Y]\{Y - \mathrm{E}[Y]\} \\
&= 2\mu(\mathbf{X})Y - \mu(\mathbf{X})^2 - \mathrm{E}[\mu(\mathbf{X})^2] - 2\mathrm{E}[Y]\{Y - \mathrm{E}[Y]\} \\
&= 2Y(\mu(\mathbf{X}) - E[Y]) - \mu(\mathbf{X})^2 + \mathrm{E}[Y]^2 - (\mathrm{E}[\mu(\mathbf{X})^2] - \mathrm{E}[Y]^2) \\
&= (Y - \mathrm{E}[Y])^2 - (Y - \mu(\mathbf{X}))^2 - \mathrm{Var}(\mu(\mathbf{X})) \qquad (4)
\end{aligned}$$

Note that $\text{Var}(\mu(\mathbf{X})) = \text{Cov}(Y, \mu(\mathbf{X}))$, so $\mathbb{IF}(\text{Var}(\mu(\mathbf{X}))) = \mathbb{IF}(\text{Cov}(Y, \mu(\mathbf{X})))$.

Plugging (4) into (3), we get

$$\begin{aligned}\mathbb{IF}(\rho_1) = &\frac{1}{2}\left(\frac{\text{Var}(Y)}{\text{Var}(\mu(\mathbf{X}))}\right)^{\frac{1}{2}}\left(\frac{((Y-\text{E}[Y])^2-(Y-\mu(\mathbf{X}))^2-\text{Var}(\mu(\mathbf{X})))\text{Var}(Y)}{\text{Var}(Y)^2}\right.\\ &\left.-\frac{\text{Var}(\mu(\mathbf{X}))((Y-\text{E}[Y])^2-\text{Var}(Y))}{\text{Var}(Y)^2}\right)\\ = &\frac{(Y-\text{E}[Y])^2-(Y-\mu(\mathbf{X}))^2}{2\rho_1\text{Var}(Y)}-\frac{\rho_1(Y-\text{E}[Y])^2}{2\text{Var}(Y)}\end{aligned}$$

#### S2.2 Mixed Influence Function

Alternatively, we can formulate the influence function for MAPA using the empirical components for terms that only involve $Y$, since we assume these can be estimated at the $\sqrt{n}$-rate. This will yield slightly different results if we start from the unsimplified or simplified parameter. We will start from the simplified version of MAPA and derive the "mixed" (subscript $M$) influence function:

$$\begin{aligned}\mathbb{IF}_M(\rho_1) &= \frac{\mathbb{IF}(\sqrt{\text{Var}(\mu(\mathbf{X}))})}{\sqrt{\text{Var}(Y)}}\\ &= \frac{\frac{1}{2}\text{Var}(\mu(\mathbf{X}))^{-\frac{1}{2}}\mathbb{IF}(\text{Var}(\mu(\mathbf{X})))}{\sqrt{\text{Var}(Y)}}\\ &= \frac{(Y-\text{E}[Y])^2-(Y-\mu(\mathbf{X}))^2-\text{Var}(\mu(\mathbf{X}))}{2\sqrt{\text{Var}(\mu(\mathbf{X}))\text{Var}(Y)}}\end{aligned}$$

#### S2.3 Parameter transformations

We compared several transformations of MAPA to improve small sample performance.

##### S2.3.1 Squared logit transformation

To ensure the one-step estimator respects the parameter bounds and to address finite sample performance, we derive the IFs for multiple transformed versions of MAPA. The squared logit transformation has the best finite sample performance of those we consider:

$$\mathbb{IF}(\text{logit}(\rho_1^2)) = \frac{2}{\rho_1(1-\rho_1^2)}\mathbb{IF}(\rho_1)$$

This transformation is defined as long as $\rho_1 \neq 0, 1$.

##### S2.3.2 Logit transformation

We also considered a logit transformation:

$$\mathbb{IF}(\text{logit}(\rho_1)) = \frac{1}{\rho_1(1-\rho_1)}\mathbb{IF}(\rho_1)$$

This transformation is defined as long as $\rho_1 \neq 0, 1$.

| Method | Acronymn | Plug-in | IF Method | Transformation |
|---|---|---|---|---|
| **Pearson Full** | PF | Pearson | Full | None |
| **Pearson Mixed** | PM | Pearson | Mixed | None |
| **Simplified Full** | SF | Simplified | Full | None |
| **Simplified Mixed** | SM | Simplified | Mixed | None |
| **Pearson Logit** | PL | Pearson | Full | Logit |
| **Simplified Logit** | SL | Simplified | Full | Logit |
| **Pearson Sq. Logit** | PQ | Pearson | Full | Squared Logit |
| **Simplified Sq. Logit** | SQ | Simplified | Full | Squared Logit |
| **Pearson Fisher** | PZ | Pearson | Full | Fisher |
| **Simplified Fisher** | SZ | Simplified | Full | Fisher |

**Table S1:** Different forms of the one-step estimator

###### S2.3.3 Fisher transformation

To compare to the standard transformation applied to Pearson correlation estimates, we also considered a Fisher transformation of the parameter:

$$\mathbb{IF}(\text{atanh}(\rho_1)) = \frac{1}{1-\rho_1^2}\mathbb{IF}(\rho_1)$$

#### S3 Constructing one-step estimators

We construct semiparametric one-step estimators by adding the empirical mean of an influence function to a plug-in estimate of the target parameter[1–6]. If we are using a transformed parameter, we transform the plug-in estimate and add the IF mean on the transformed scale. For MAPA, there are two potential plug-in estimates. The first is the standard Pearson estimate,

$$\hat{\rho}_P = \frac{\widehat{\text{Cov}}(Y, \hat{\mu}(\mathbf{X}))}{\sqrt{\widehat{\text{Var}}(Y)\widehat{\text{Var}}(\hat{\mu}(\mathbf{X}))}}.$$

The second option is to use the empirical variance estimates for the simplified version of MAPA:

$$\hat{\rho}_S = \frac{\sqrt{\widehat{\text{Var}}(\hat{\mu}(\mathbf{X}))}}{\sqrt{\widehat{\text{Var}}(Y)}}$$

**Table S1** summarizes all estimators, and the full formulas for each are in Section S11.

##### S3.1 Evaluating parameter transformations

To select among the possible transformation methods, we evaluated all transformations in the Adolescent Brain Cognitive Development (ABCD) study [7]. We used the ABCD so that we did not select and evaluate a method in the same dataset. For this analysis we used the same simulation procedure with age as the phenotype and harmonized structural data averaged within the Desikan-Killiany atlas as the features (**Figure S1**).

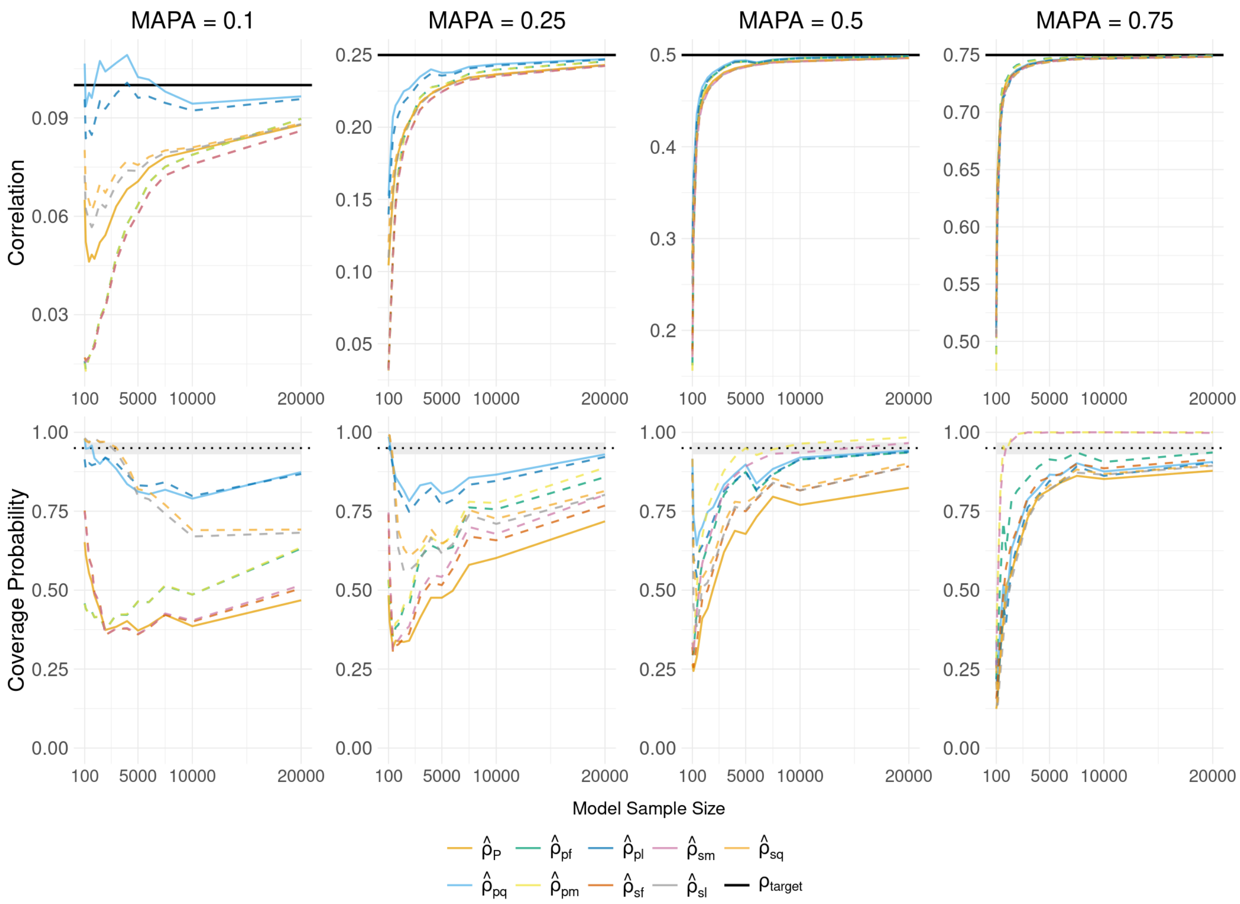


**Figure S1: The squared-logit transformation (PQ) has the smallest bias across sample sizes and values of MAPA in the ABCD. A)** Mean estimates of the Pearson (solid orange) and one-step estimators based on different parameter transformations and influence function construction methods. The data is harmonized structural data (DK atlas, 68 regions) from ABCD, using ridge regression to predict age. **B)** 95% confidence interval coverage probability for the Pearson estimator and each version of the one-step estimator. The PQ estimator (solid blue) has the best coverage for large sample sizes and small values of MAPA and the most consistent performance across values of MAPA. For the estimator acronyms, the first letter indicates whether the plug-in estimator used is Pearson (P) or the simplified version (S). The second letter denotes the transformation: F = untransformed, M = Pearson untransformed, using empirical components of $Y$ where possible when constructing influence function, L = logit transformation, Q = squared logit transformation.

### S4 Requirements for asymptotic normality

Semiparametric and empirical process theory give conditions for the one-step estimator to achieve asymptotic normality[1–6]. Without using sample splitting, we must assume $\mu$ is not overly complex. For example, we could restrict to a Donsker class[5,6]. This can be too restrictive for many modern ML methods. Using sample splitting and cross-fitting removes the need to employ a Donsker-type restriction. For the one-step estimator to converge to a normal distribution at the $\sqrt{n}$-rate when applying cross-fitting, $\hat{\mu}$ must converge to $\mu$ in $L2$ norm at a rate of $n^{-\frac{1}{4}}$ or faster[5].

We evaluated the convergence rates for the Pearson and one-step estimator settings for the structural data predicting age (**Figure 2**) for various values of MAPA using non-linear least squares and the power law formula

$$\mathbb{E}\hat{\rho} = \alpha n^{\beta} + \rho_1. \tag{5}$$

For MAPA of 0.25, 0.5, and 0.75 with random forest, Pearson's correlation estimator is estimated to converge at a rate between $n^{-1/2}$ and $n^{-1/4}$, and confidence intervals fail (**Table S2**).

| **ML Model** | MAPA | **Pearson Convergence** | **One-Step Convergence** |
|---|---|---|---|
| Ridge | 0.1 | -0.19 (-0.21, -0.17) | -0.25 (-0.30, -0.20) |
| Ridge | 0.25 | -0.46 (-0.48, -0.44) | -1.06 (-1.19, -0.95) |
| Ridge | 0.5 | -0.64 (-0.66, -0.62) | -0.91 (-0.95, -0.87) |
| Ridge | 0.75 | -0.80 (-0.82, -0.78) | -0.97 (-1.00, -0.94) |
| Random Forest | 0.1 | -0.15 (-0.17, 0.13) | |
| Random Forest | 0.25 | -0.34 (-0.36, -0.33) | -0.41 (-0.45, -0.37) |
| Random Forest | 0.5 | -0.37 (-0.38, -0.35) | -0.60 (-0.62, -0.58) |
| Random Forest | 0.75 | -0.33 (-0.33, -0.32) | -0.66 (-0.68, -0.65) |

**Table S2: The one-step estimator has a faster estimated convergence rate ($\beta$ in Eqn.** (5)**) to MAPA than the Pearson estimator**. For the one-step estimator, convergence is faster, and near-nominal coverage is retained across settings. The non-linear least squares model did not converge for MAPA = 0.1 for random forest.

To empirically assess the standard error estimation of the one-step estimator, we plot mean estimated standard error against the true standard deviation across the simulations for predicting age from structural data (**Figure S2**). The standard error is underestimated for small samples but asymptotically close to the true standard deviation. The small-sample underestimation is less severe for higher values of MAPA.

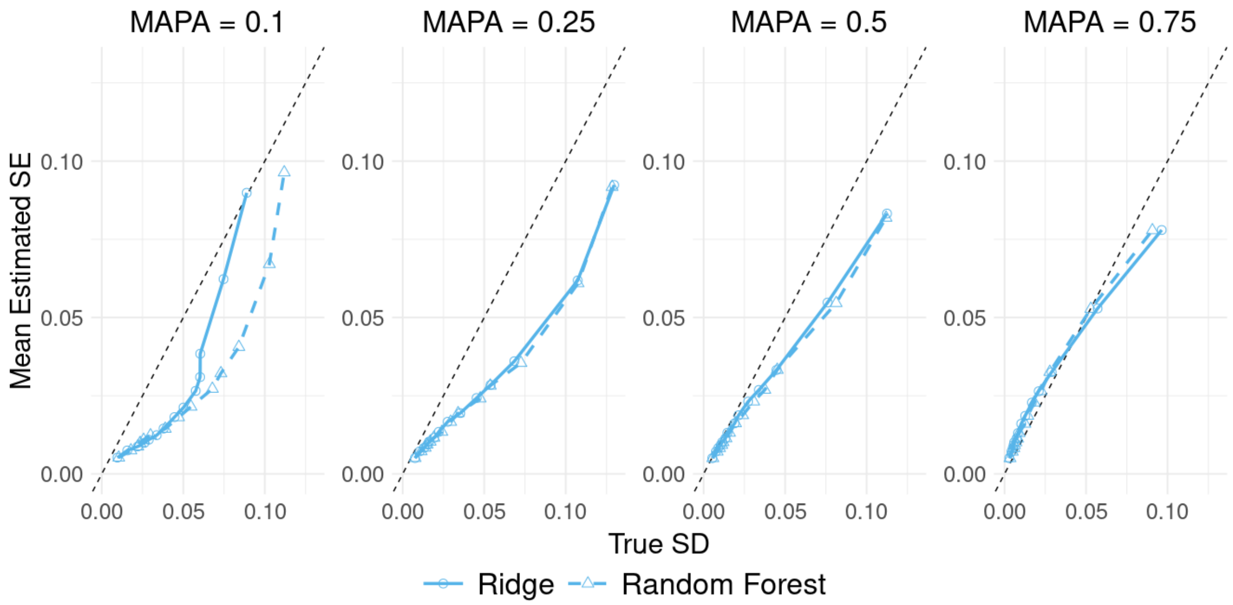


**Figure S2: The estimated one-step standard error is asymptotically close to the true standard deviation of the estimator across simulations.** The mean standard error estimate (y-axis) is asymptotically close to the true standard deviation of the one-step estimator across 500 simulations (x-axis) for both ridge regression (solid lines, circles) and random forest (dashed lines, triangles) models.

### S5 Methods to combine estimators across folds

Because we perform cross-fitting to relax complexity constraints on $\hat{\mu}(\mathbf{X})$, we must specify how to aggregate the resulting fold-specific estimates. Zhou et al.[8] describes two approaches to averaging correlation estimates from a cross-fitting setting: "Instant", where the correlation point estimate is calculated for each fold and the resulting $K$ estimates are averaged for the the final point estimate, and "Hold", where the needed components of the correlation estimate are pooled across all folds to compute one point estimate at the end. Typical semiparametric estimators follow the "instant" approach, but we additionally assessed performance for a "hold" approach (**Figure S3**) for comparison.

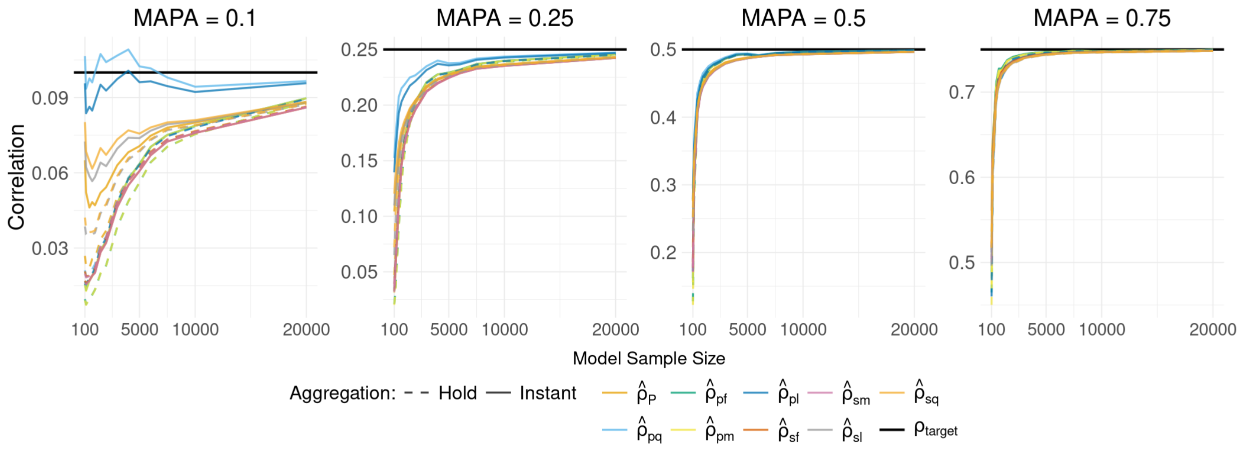


**Figure S3: Averaging fold-specific correlation estimates outperforms pooling observations from all folds in the ABCD.** Mean estimates of the Pearson (orange) and one-step estimators based on different parameter transformations for different strategies for aggregating estimates across folds. The "instant" method (solid lines) computes one-step estimates in each fold from fold-specific influence function means and averages the resulting estimates across folds for a final estimator. This method has smaller bias across estimator types than the "hold" method, which pools all influence function estimates together to construct a final one-step estimator using all folds. These simulations used harmonized structural data from the ABCD, using ridge regression to predict age.

### S6 Additional simulation results

#### S6.1 Evaluation of dimensionality with functional connectivity data

We varied the resolution of the functional connectivity (FC) networks in the simulations to evaluate the effect of data dimensionality on convergence rates of the correlation estimators when using ridge regression. To do this, FC was computed in the Schaefer parcellation and averaged at 7-network or 17-Yeo network resolutions (**Figure S4**).

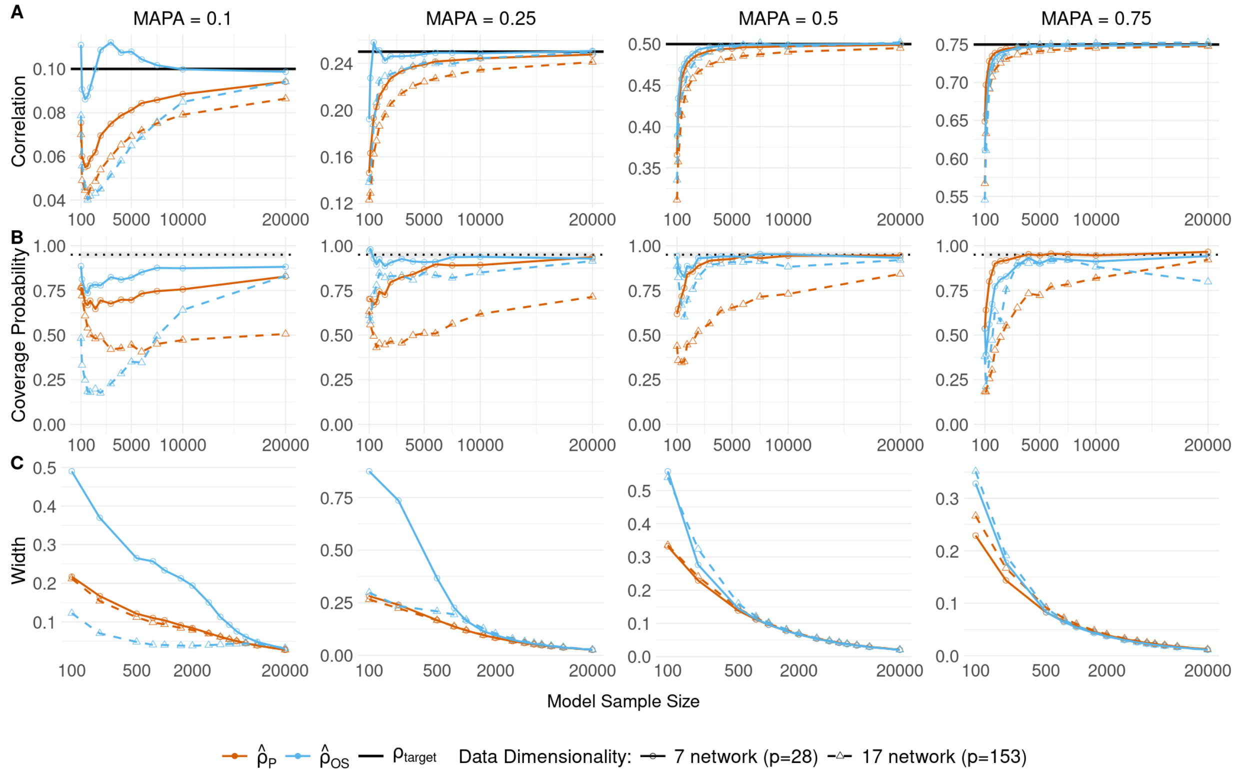


**Figure S4: The one-step estimator outperforms Pearson's estimator for most values of MAPA across data dimensionalities in plasmode simulations of FC with age using ridge regression models. A)** Mean estimates of the Pearson (orange) and one-step (blue) correlation for ridge regression models using either 7 network FC data (solid line, circles; number of predictors = 28) or 17 network FC data (dashed line, triangles; number of predictors = 153) for different values of MAPA. **B)** 95% confidence interval coverage probability using the Fisher interval (for Pearson) or the one-step confidence interval. The gray shaded region is the expected range of coverage for 500 simulations. **C)** Average confidence interval width for each method.

#### S6.2 Evaluation of psychiatric prediction with structural data

Our main simulations predicted age in the build model in order to generate simulated phenotypes. We used also simulations to evaluate the effect of data dimensionality on convergence rates when predicting simulated P-factor by using the gray matter volume (GMV) averaged in the Desikan-Killiany-Tourville atlas (62 features) and GMV in the Schaefer 300 atlas (**Figure S5**).

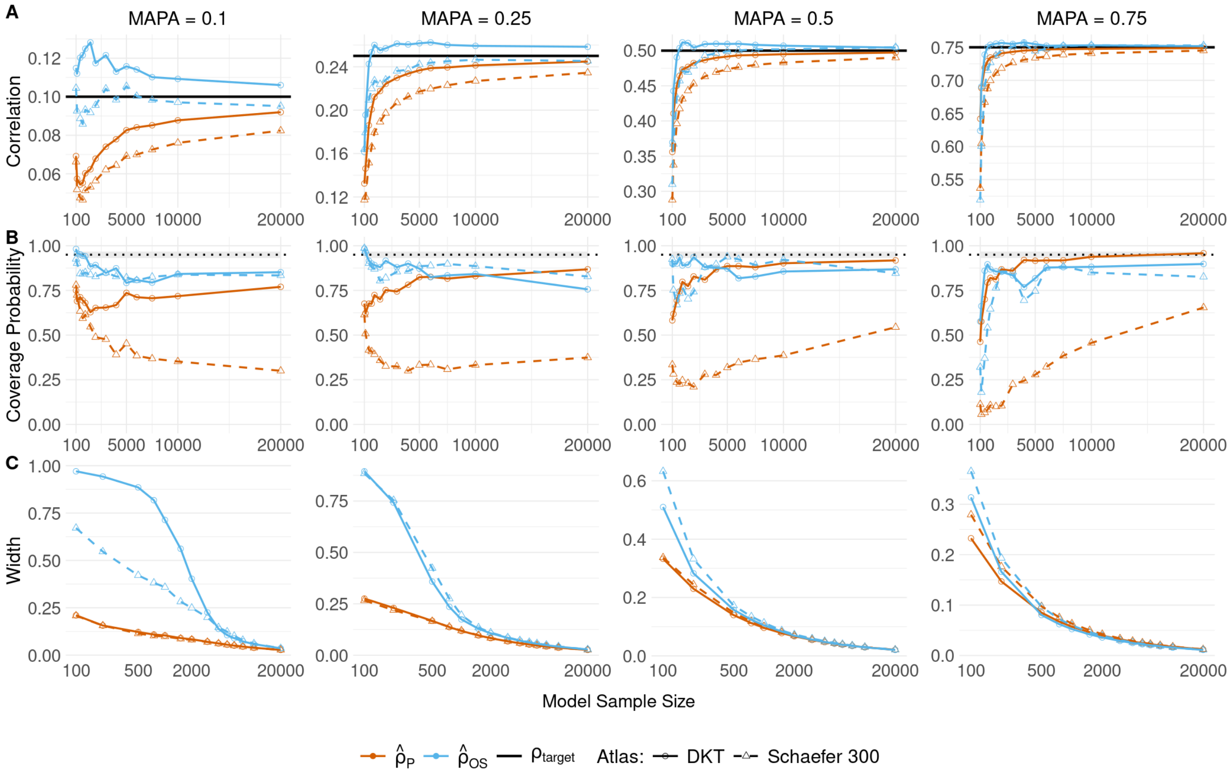


**Figure S5: The one-step estimator outperforms Pearson's estimator across data dimensionalities in realistic plasmode simulations of anatomical (62 features) association with P-Factor. A)** Mean estimates of the Pearson (orange) and one-step (blue) correlation across 500 simulations of ridge regression models predicting P-Factor using structural data from the DKT atlas (solid line, circles) and Schaefer 300 atlas (dashed line, triangles) models for different values of MAPA. **B)** 95% confidence interval coverage probability using the Fisher interval (for Pearson) or the one-step confidence interval. The gray shaded region is the 95% probability interval for the 500 simulations. **C)** Average confidence interval width for each method.

### S7 Comparing models via the Delta Method

We can leverage the asymptotic normality of the semiparametric one-step estimator to construct estimators with confidence intervals for comparing MAPA for multiple model classes (e.g., comparing two ML algorithms or two sets of features). Denote MAPA with $\rho$. Let $\hat{\rho}_1$ and $\hat{\rho}_2$ denote the one-step estimates for $\rho_1$ and $\rho_2$ for two different model classes. We know

$$\sqrt{n}\left\{\begin{pmatrix}\hat{\rho}_1\\ \hat{\rho}_2\end{pmatrix} - \begin{pmatrix}\rho_1\\ \rho_2\end{pmatrix}\right\} \xrightarrow{d} \mathcal{N}\left(\begin{pmatrix}0\\ 0\end{pmatrix}, \mathbf{\Sigma}\right)$$

where $\mathbf{\Sigma} = \begin{pmatrix}\mathrm{Var}(\mathbb{IF}(\rho_1)) & \mathrm{Cov}(\mathbb{IF}(\rho_1), \mathbb{IF}(\rho_2))\\ \mathrm{Cov}(\mathbb{IF}(\rho_1), \mathbb{IF}(\rho_2)) & \mathrm{Var}(\mathbb{IF}(\rho_2))\end{pmatrix}$.

Note, we present one-step estimators based on parameter transformations. Consider a transformation $t(\rho)$ with estimate $\hat{\rho}_t$ (i.e., the one-step estimate before back-transforming to the correlation scale)

$$\sqrt{n}\left\{\begin{pmatrix}\hat{\rho}_{t,1}\\ \hat{\rho}_{t,2}\end{pmatrix} - \begin{pmatrix}t(\rho_1)\\ t(\rho_2)\end{pmatrix}\right\} \xrightarrow{d} \mathcal{N}\left(\begin{pmatrix}0\\ 0\end{pmatrix}, \mathbf{\Sigma}_t\right)$$

where $\mathbf{\Sigma}_t = \begin{pmatrix} \text{Var}(\mathbb{IF}(t(\rho_1))) & \text{Cov}(\mathbb{IF}(t(\rho_1)), \mathbb{IF}(t(\rho_2))) \\ \text{Cov}(\mathbb{IF}(t(\rho_1)), \mathbb{IF}(\rho_2)) & \text{Var}(\mathbb{IF}(t(\rho_2))) \end{pmatrix}$.

#### S7.1 Comparing the difference in correlations

To compare $\rho_2 - \rho_1$ where $\hat{\rho} = t^{-1}(\hat{\rho}_t)$ we use the multivariate delta method:

$$\begin{aligned} g(t(\rho)) &= t^{-1}(t(\rho_2)) - t^{-1}(t(\rho_1)) \\ &= \rho_2 - \rho_1 \end{aligned}$$

$$\sqrt{n}\left\{ g\begin{pmatrix} \hat{\rho}_{t,1} \\ \hat{\rho}_{t,2} \end{pmatrix} - g\begin{pmatrix} t(\rho_1) \\ t(\rho_2) \end{pmatrix} \right\} \xrightarrow{d} \mathcal{N}\left(0, g'(t(\rho))^T \mathbf{\Sigma}_t g(t(\rho))\right)$$

where $g'(t(\rho)) = \begin{pmatrix} -(t^{-1})'(\hat{\rho}_{t,1}) \\ (t^{-1})'(\hat{\rho}_{t,2}) \end{pmatrix}$.

For $t(\rho) = \text{logit}(\rho^2)$, $(t^{-1})'(\hat{\rho}_t) = \frac{1}{2}\hat{\rho}_t(1-\hat{\rho}_t^2)$. This yields the variance estimate for the confidence interval

$$\begin{aligned} &\frac{1}{4}\widehat{\text{Var}}\left\{\hat{\mathbb{IF}}(t(\rho_1))\right\} \hat{\rho}_1^2(1-\hat{\rho}_1^2)^2 \\ &\quad + \frac{1}{4}\widehat{\text{Var}}\left\{\hat{\mathbb{IF}}(t(\rho_2))\right\} \hat{\rho}_2^2(1-\hat{\rho}_2^2)^2 \\ &\quad - \frac{1}{2}\widehat{\text{Cov}}\left\{\hat{\mathbb{IF}}(t(\rho_1)), \hat{\mathbb{IF}}(t(\rho_2))\right\} \hat{\rho}_1\hat{\rho}_2(1-\hat{\rho}_1^2)(1-\hat{\rho}_2^2) \end{aligned}$$

.

Because MAPA is bounded in [0,1], we truncate confidence intervals for the difference at -1 and 1.

#### S7.2 Comparing the effect size ratio

To compare correlation values from two models, taking the difference on the squared logit corresponds to taking the ratio of Cohen's $f^2 = \frac{\rho^2}{1-\rho^2}$ for the two models[9]. Thus, to get a confidence interval for the ratio of the effect sizes for two different model classes, we can take a difference on the transformed scale.

$$\begin{aligned} \text{logit}(\rho_2^2) - \text{logit}(\rho_1^2) &= \log(f_2^2) - \log(f_1^2) \\ &= \log\left(\frac{f_2^2}{f_1^2}\right). \end{aligned}$$

We construct confidence intervals as

$$\begin{aligned} &\text{logit}(\hat{\rho}_2^2) - \text{logit}(\hat{\rho}_1^2) \pm \\ &\frac{z_{1-\alpha/2}}{\sqrt{n}}\sqrt{\widehat{\text{Var}}\big(\hat{\mathbb{IF}}\{\text{logit}(\rho_1^2)\}\big) + \widehat{\text{Var}}\big(\hat{\mathbb{IF}}\{\text{logit}(\rho_2^2)\}\big) - 2\,\widehat{\text{Cov}}\big(\hat{\mathbb{IF}}\{\text{logit}(\rho_1^2)\}, \hat{\mathbb{IF}}\{\text{logit}(\rho_2^2)\}\big)} \end{aligned}$$

to get a log-ratio of the variance explanation.

We applied this method in the RBC to visualize the ratio of the variance explained among the different model types (**Figure S6**).

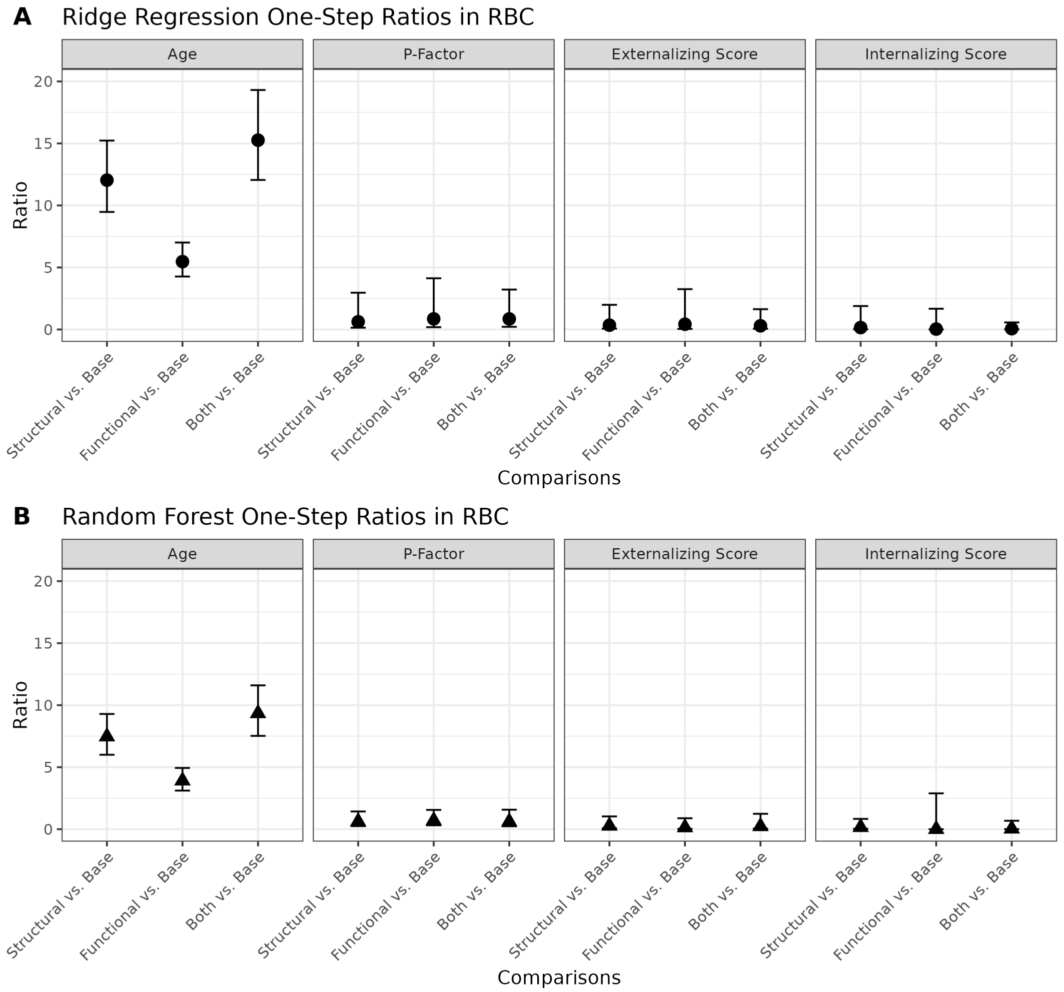


**Figure S6: The ratio of one-step estimators indicates increased prediction accuracy for age using imaging data.** Median estimate across 250 sample splits comparing prediction accuracy via a ratio with confidence intervals constructed with the delta method between structural, functional, or both versus no imaging data ("Base") for the one-step estimator using (**A**) random forest and (**B**) ridge regression for four outcome phenotypes.

### S8 Simulation error distribution

In each simulation replicate, we resample rows from the evaluation dataset (i.e., the 80% of the original dataset not used to train the build model), including the input features and the "true" conditional mean computed as the prediction from the build model. We then generate a simulated value for the phenotype by adding a random noise term $\epsilon$ from a uniform distribution, with bounds chosen to control the variance and thus MPA.

$$\rho_0 = \sqrt{\frac{\text{Var}(\xi(\mathbf{X}))}{\text{Var}(Y)}}$$

$$= \sqrt{\frac{\text{Var}(\xi(\mathbf{X}))}{\text{Var}(\xi(\mathbf{X})) + \text{Var}(\epsilon)}}.$$

$$\text{Var}(\epsilon) = \text{Var}(\xi(\mathbf{X})) \times \frac{1-\rho_0^2}{\rho_0^2}$$

For the uniform distribution, this yields the bounds:

$$\epsilon \sim \text{Unif}(-a, a);$$

$$a = \sqrt{3 \times \text{Var}(\xi(\mathbf{X})) \times \frac{1-\rho_0^2}{\rho_0^2}}$$

### S9 Computing ground truth TPA

To compute the ground truth TPA for the simulation in Figure 1, we trained a ridge regression model $\hat{\mu}$ in 500 samples of the evaluation dataset, with the phenotype generated as described above. Then, in each resample of the remaining (i.e., testing) data, we computed the correlation between the $\hat{\mu}$ predictions and the true conditional mean $\xi$, adjusted for variability in the phenotype, with the formula below

$$\text{TPA} = \frac{\hat{\text{E}}[\hat{\mu}(\mathbf{X})\xi(\mathbf{X})] - \hat{\text{E}}[\hat{\mu}(\mathbf{X})]\hat{\text{E}}[\xi(\mathbf{X})]}{\sqrt{\widehat{\text{Var}}(\hat{\mu}(\mathbf{X}))(\widehat{\text{Var}}(\xi(\mathbf{X})) + \text{Var}(\epsilon))}}$$

with the error variance computed as described above. We defined the ground truth TPA as the average over 500 simulations at the largest simulated sample size, $n$ = 20,000.

### S10 Single-sample evaluation in HBN Staten Island site

We performed analyses in the Staten Island site of the Healthy Brain Network (HBN) data to evaluate performance of the one-step estimator in a smaller dataset [10] ($n$ = 208, **Table S3**). Compared to the analysis in with the full dataset ($n$ = 3074), the confidence intervals are wider **Figure S7**. For the psychiatric phenotypes, the random forest models and baseline ridge models have especially wide confidence intervals, highlighting issues near the boundary (i.e., 0) of the one-step estimator. Theoretically, the one-step estimator we used assumes the parameter is not exactly equal to 0, and as the true value gets very near zero, the estimated variance can become unstable. Similar to the full dataset analysis, there is an increase in predictive accuracy for adding structural or structural plus functional imaging to predict age, but not the psychiatric phenotypes **Figure S8**.

| | Overall (n=208) |
|---|---|
| **Age (years)** | |
| Mean (SD) | 11.7 (3.35) |
| Median [Min, Max] | 11.0 [5.40, 18.0] |
| **Sex** | |
| Female | 88 (42.3%) |
| Male | 120 (57.7%) |
| **Internalizing Score** | |
| Mean (SD) | -0.0226 (1.05) |
| Median [Min, Max] | -0.201 [-2.17, 2.82] |
| **Externalizing Score** | |
| Mean (SD) | 0.0319 (1.01) |
| Median [Min, Max] | -0.243 [-2.15, 3.89] |
| **P-Factor** | |
| Mean (SD) | 0.181 (1.02) |
| Median [Min, Max] | 0.259 [-1.61, 2.77] |
| **Mean Framewise Displacement** | |
| Mean (SD) | 0.132 (0.138) |
| Median [Min, Max] | 0.0773 [0.0294, 0.837] |
| **Euler Number** | |
| Mean (SD) | -66.6 (33.4) |
| Median [Min, Max] | -58.0 [-232, -17.0] |

**Table S3: Participant characteristics in the Healthy Brain Network (HBN) Staten Island site, used for supplementary analyses.**

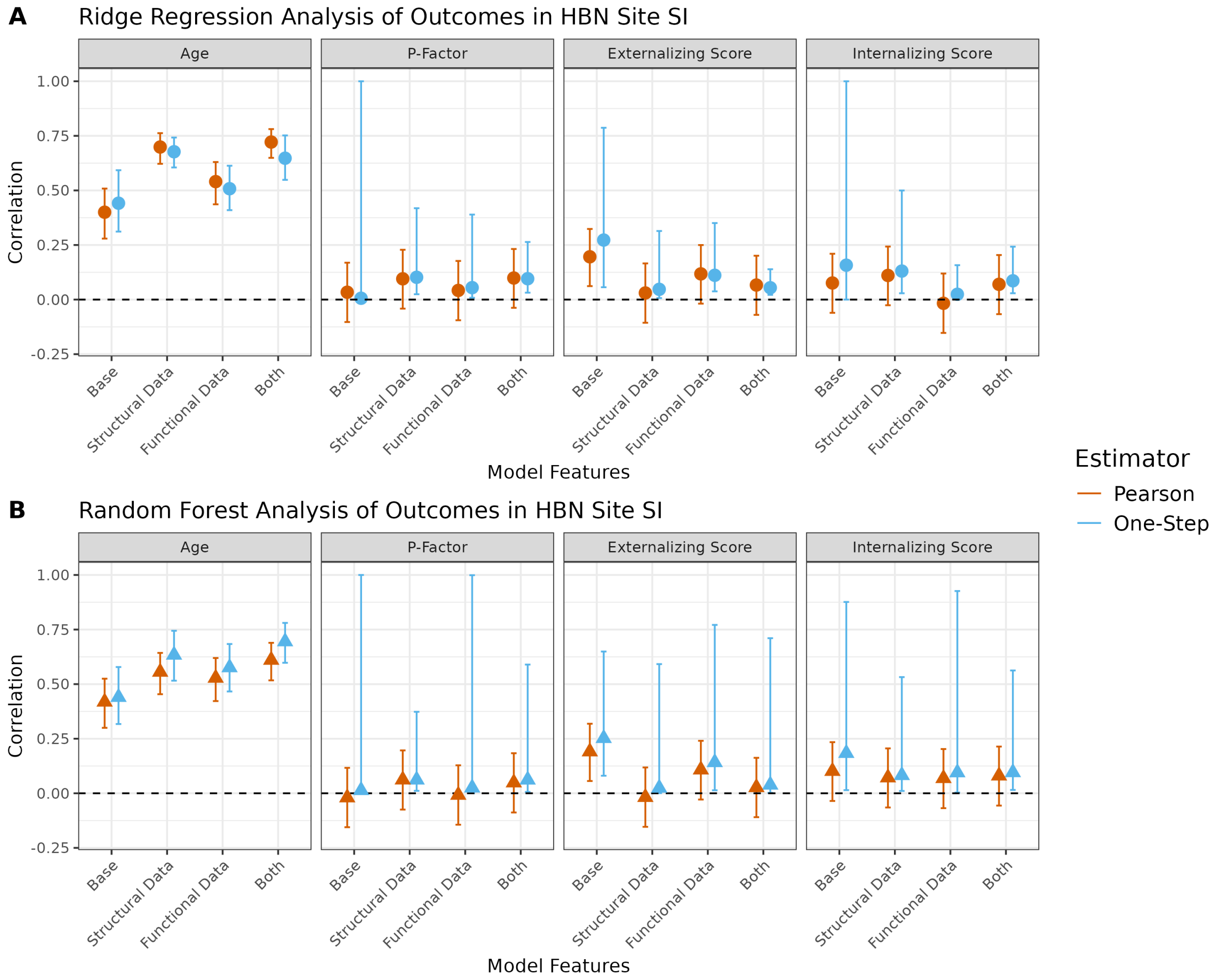


**Figure S7: Confidence intervals are wider for small sample sizes and low predictive accuracy.** Median estimates across 250 sample splits of the HBN Staten Island data (n = 208) of the Pearson and one-step correlation for (A) random forest models and (B) ridge regression using no imaging data ("Base"), structural data, functional data, or both as model features predicting four outcome phenotypes, shown with 95% confidence intervals. For most associations, the one-step estimator has a slightly larger point estimate. The confidence intervals are much wider for the one-step estimators for the psychopathologic phenotypes, which have lower point estimates compared to age.

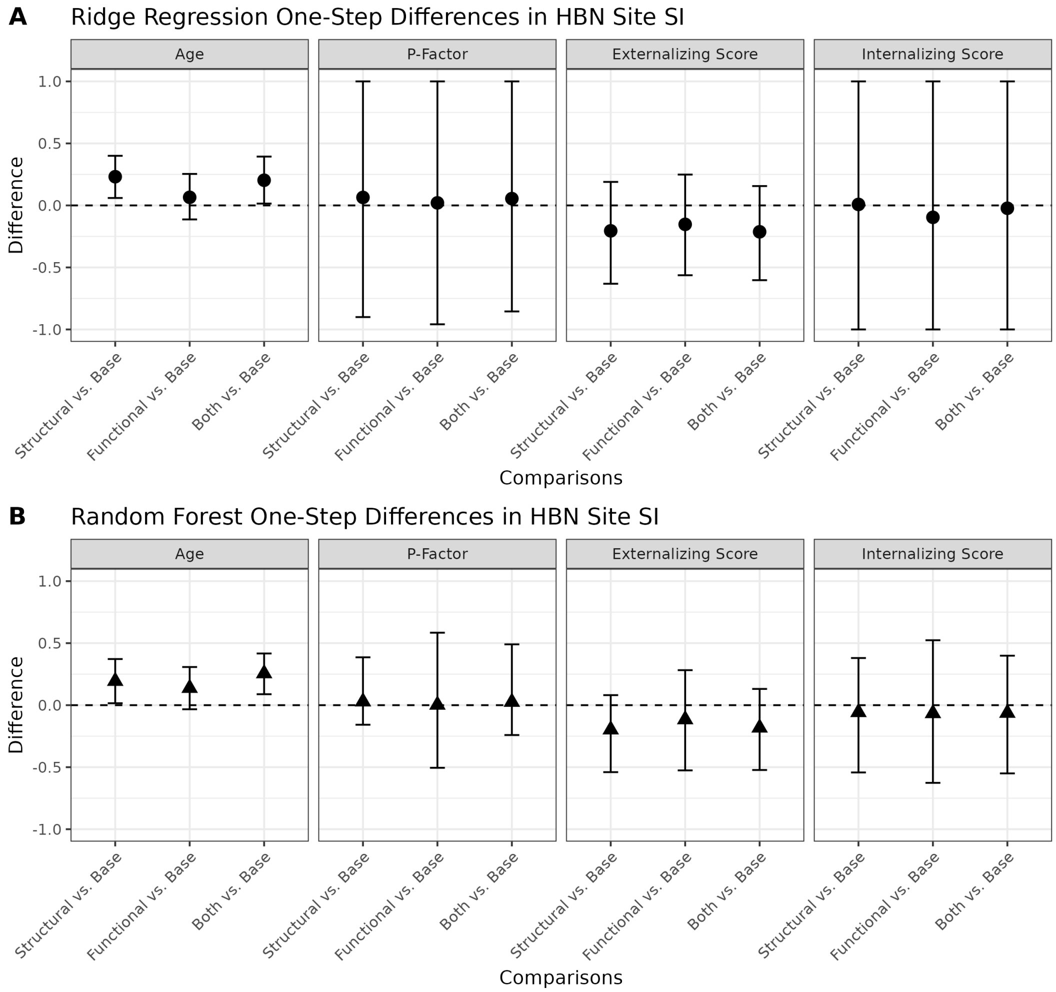


**Figure S8: Improvement of prediction accuracy for age using imaging data is captured in a smaller sample.** Median estimate across 250 sample splits of the HBN Staten Island data (n = 208) comparing prediction accuracy between structural, functional, or both versus no imaging data ("Base") for the one-step estimator using (A) random forest and (B) ridge regression for four outcome phenotypes. For age, the structural and combination imaging data improve prediction accuracy.

### S11 One-step estimator formulas

Note: $\mathbb{P}_n$ indicates the sample average.

- "Pearson Full (PF)":

$$\hat{\rho}_{PF} = \hat{\rho}_P + \mathbb{P}_n \left\{ \frac{(Y - \hat{\mathrm{E}}[Y])^2 - (Y - \hat{\mu}(\mathbf{X}))^2}{2\sqrt{\widehat{\mathrm{Var}}(\hat{\mu}(\mathbf{X}))\widehat{\mathrm{Var}}(Y)}} - \frac{\sqrt{\widehat{\mathrm{Var}}(\hat{\mu}(\mathbf{X}))}(Y - \hat{\mathrm{E}}[Y])^2}{2\widehat{\mathrm{Var}}(Y)^{\frac{3}{2}}} \right\}$$

- "Pearson Mixed (PM)":

$$\hat{\rho}_{PM} = \hat{\rho}_P + \mathbb{P}_n \left\{ \frac{(Y - \hat{\mathrm{E}}[Y])^2 - (Y - \hat{\mu}(\mathbf{X}))^2 - \widehat{\mathrm{Var}}(\hat{\mu}(\mathbf{X}))}{2\sqrt{\widehat{\mathrm{Var}}(\hat{\mu}(\mathbf{X}))\widehat{\mathrm{Var}}(Y)}} \right\}$$

- "Simplified Full (SF)":

$$\hat{\rho}_{SF} = \hat{\rho}_S + \mathbb{P}_n \left\{ \frac{(Y - \hat{\mathrm{E}}[Y])^2 - (Y - \hat{\mu}(\mathbf{X}))^2}{2\sqrt{\widehat{\mathrm{Var}}(\hat{\mu}(\mathbf{X}))\widehat{\mathrm{Var}}(Y)}} - \frac{\sqrt{\widehat{\mathrm{Var}}(\hat{\mu}(\mathbf{X}))}(Y - \hat{\mathrm{E}}[Y])^2}{2\widehat{\mathrm{Var}}(Y)^{\frac{3}{2}}} \right\}$$

- "Simplified Mixed (SM)":

$$\hat{\rho}_{SM} = \hat{\rho}_S + \mathbb{P}_n \left\{ \frac{(Y - \hat{\mathrm{E}}[Y])^2 - (Y - \hat{\mu}(\mathbf{X}))^2 - \widehat{\mathrm{Var}}(\hat{\mu}(\mathbf{X}))}{2\sqrt{\widehat{\mathrm{Var}}(\hat{\mu}(\mathbf{X}))\widehat{\mathrm{Var}}(Y)}} \right\}$$

- "Pearson Logit (PL)":

$$\hat{\rho}_{PL} = \mathrm{expit}\Bigg( \mathrm{logit}(\hat{\rho}_P) + \mathbb{P}_n \Bigg\{ \Big( \frac{(Y - \hat{\mathrm{E}}[Y])^2 - (Y - \hat{\mu}(\mathbf{X}))^2}{2\,\widehat{\mathrm{Var}}(\hat{\mu}(\mathbf{X}))} - \frac{(Y - \hat{\mathrm{E}}[Y])^2}{\widehat{\mathrm{Var}}(Y)} \Big) \times \Big( \frac{\sqrt{\widehat{\mathrm{Var}}(Y)}}{\sqrt{\widehat{\mathrm{Var}}(Y)} - \sqrt{\widehat{\mathrm{Var}}(\hat{\mu}(\mathbf{X}))}} \Big) \Bigg\} \Bigg)$$

- "Simplified Logit (SL)":

$$\hat{\rho}_{SL} = \mathrm{expit}\Bigg( \mathrm{logit}(\hat{\rho}_S) + \mathbb{P}_n \Bigg\{ \Big( \frac{(Y - \hat{\mathrm{E}}[Y])^2 - (Y - \hat{\mu}(\mathbf{X}))^2}{2\,\widehat{\mathrm{Var}}(\hat{\mu}(\mathbf{X}))} - \frac{(Y - \hat{\mathrm{E}}[Y])^2}{\widehat{\mathrm{Var}}(Y)} \Big) \times \Big( \frac{\sqrt{\widehat{\mathrm{Var}}(Y)}}{\sqrt{\widehat{\mathrm{Var}}(Y)} - \sqrt{\widehat{\mathrm{Var}}(\hat{\mu}(\mathbf{X}))}} \Big) \Bigg\} \Bigg)$$

- "Pearson Squared Logit (PQ)":

$$\hat{\rho}_{PQ} = \sqrt{ \mathrm{expit}\left( \mathrm{logit}(\hat{\rho}_P^2) + \mathbb{P}_n \left\{ \frac{(Y - \mathrm{E}[Y])^2 - \widehat{\mathrm{Var}}(Y)(Y - \hat{\mu}(\mathbf{X}))^2}{\widehat{\mathrm{Var}}(\hat{\mu}(\mathbf{X}))(\widehat{\mathrm{Var}}(Y) - \widehat{\mathrm{Var}}(\hat{\mu}(\mathbf{X})))} \right\} \right) }$$

- "Simplified Squared Logit (SQ)":

$$\hat{\rho}_{SQ} = \sqrt{ \mathrm{expit}\left( \mathrm{logit}(\hat{\rho}_S^2) + \mathbb{P}_n \left\{ \frac{(Y - \mathrm{E}[Y])^2 - \widehat{\mathrm{Var}}(Y)(Y - \hat{\mu}(\mathbf{X}))^2}{\widehat{\mathrm{Var}}(\hat{\mu}(\mathbf{X}))(\widehat{\mathrm{Var}}(Y) - \widehat{\mathrm{Var}}(\hat{\mu}(\mathbf{X})))} \right\} \right) }$$

- "Pearson Fisher (PZ)":

$$\hat{\rho}_{PZ} = \tanh\left( \mathrm{atanh}(\hat{\rho}_P) + \mathbb{P}_n \left\{ \frac{(Y - \hat{\mathrm{E}}[Y])^2 - (Y - \hat{\mu}(\mathbf{X}))^2}{2(1 - \hat{\rho}_P^2)\sqrt{\widehat{\mathrm{Var}}(\hat{\mu}(\mathbf{X}))\widehat{\mathrm{Var}}(Y)}} - \frac{\sqrt{\widehat{\mathrm{Var}}(\hat{\mu}(\mathbf{X}))}(Y - \hat{\mathrm{E}}[Y])^2}{2(1 - \hat{\rho}_P^2)\widehat{\mathrm{Var}}(Y)^{\frac{3}{2}}} \right\} \right)$$

- "Simplified Fisher (SZ)":

$$\hat{\rho}_{SZ} = \tanh\left( \mathrm{atanh}(\hat{\rho}_S) + \mathbb{P}_n \left\{ \frac{(Y - \hat{\mathrm{E}}[Y])^2 - (Y - \hat{\mu}(\mathbf{X}))^2}{2(1 - \hat{\rho}_P^2)\sqrt{\widehat{\mathrm{Var}}(\hat{\mu}(\mathbf{X}))\widehat{\mathrm{Var}}(Y)}} - \frac{\sqrt{\widehat{\mathrm{Var}}(\hat{\mu}(\mathbf{X}))}(Y - \hat{\mathrm{E}}[Y])^2}{2(1 - \hat{\rho}_P^2)\widehat{\mathrm{Var}}(Y)^{\frac{3}{2}}} \right\} \right)$$
